# Myeloid cell evolution uncovered by shrimp immune cell analysis at single-cell resolution

**DOI:** 10.1101/2022.05.17.492277

**Authors:** Peng Yang, Yaohui Chen, Zhiqi Huang, Huidan Xia, Ling Cheng, Hao Wu, Yueling Zhang, Fan Wang

## Abstract

How myeloid cells evolved from invertebrate to vertebrate is still a mystery. Here we collected circulating hemocytes from a marine invertebrate-*Penaeus vannamei* via gradient centrifugation and identified prohemocyte, monocytic hemocyte and granulocyte as three major types of cells in shrimp hemolymph by single-cell mRNA sequencing. Additional pseudotime trajectory analysis revealed that shrimp monocytic hemocytes and granulocytes were differentiated from a common progenitor which was similar with that of human myeloid cells. More interestingly, we identified that MH2, a terminal differentiated monocytic hemocyte, was a macrophage-like phagocytic cell which could engulf fluorescein labelled *Vibrio parahaemolyticus* and shared nine marker genes including inflammasome components *Nlrp3* and *Casp1* with human macrophage. After that, we compared our classification with traditional shrimp hemocytes classification and found that hyalinocyte included both prohemocyte and monocytic hemocyte while semi-granulocyte included both monocytic hemocyte and granulocyte. In general, our results redefine shrimp hemocyte classification based on functional marker genes and unveil evolutionary trace of myeloid cells in marine invertebrate.

## Introduction

Myeloid cells are a major component of the immune system comprising monocytes and granulocytes, in which monocytes could differentiate into dendritic cells and macrophages in different microenvironments. All these cells constitute the immediate immune defense line against various pathogens invasion(Bassler, Schulte-Schrepping, Warnat-Herresthal, Aschenbrenner, & Schultze, 2019). However, how these cells evolved is still unclear. Recently, advanced single-cell sequencing technology sheds the light to answer this question. For example, recent progress on mosquito cellular immunity unveiled that mosquito hemocyte could be divided into four major types (prohemocyte, granulocyte, oenocytoid, and megacyte)(Kwon, Mohammed, Franzen, Ankarklev, & Smith, 2021; Raddi et al., 2020). Hemocytes from another invertebrate model-*Drosophila menlangaster* could be divided into eight subgroups including crystal cells, lamellocyte, unspecified plasmatocytes, proliferative plasmatocytes, PSC-like hemocytes, antimicrobial plasmatocytes, phagocytic plasmatocytes and secretory plasmatocytes(Cattenoz, Monticelli, Pavlidaki, & Giangrande, 2021; Cho et al., 2020; H. Li et al., 2022; Tattikota et al., 2020). All these classifications cover the major innate immune cell functions including proliferation, reactive oxygen species (ROS) generation, phagocytosis, and effector secretion, which are similar with vertebrate myeloid cells. Thus, an interesting question is raised: are there any direct evidences showing that the vertebrate myeloid cells originate from invertebrate?

Intrigued by this question, we applied a marine invertebrate-*Penaeus vannamei* as a model to further explore it. *Penaeus vannamei*, a popular mariculture species featured with its fast-growing and delicious taste, has attracted more and more attention in recent years(Zhang et al., 2019). This invertebrate animal has an open circulating system filled with hemolymph which consists of plasma and hemocyte(Lin & Soderhall, 2011). The functions of shrimp hemolymph are highly similar with that of human peripheral blood including transportation of all kinds of nutrients and metabolic waste, maintenance of acid-base equilibrium, defense of various invaded pathogens and hemostatic effect(McNamara & Faria, 2012; Tassanakajon et al., 2018). Moreover, its circulating plasma contains more than 400 various proteins, many of which are human homologs(Luo, Chen, & Wang, 2022; Tao et al., 2019); its circulating hemocytes contain proliferating cells, phagocytic cells and effector secreting cells(Lin & Soderhall, 2011), which are functionally similar with human peripheral myeloid cells. With these advantages, we systemically explored shrimp plasma proteins and identified that CREG is a shrimp hemocyte activation factor(Huang, Yang, & Wang, 2021; Tao et al., 2019). In this study, we further explored CREG effect via single-cell sequencing of shrimp hemocytes and redefined shrimp hemocyte classification according to their functional marker genes distribution.

## Results

### Major cell types of shrimp circulating hemocytes

Previously we identified an LPS-induced shrimp plasma protein-CREG. Injection of recombinant CREG (rCREG) could effectively activate shrimp hemocyte compared with recombinant EGFP (rEGFP) injected group(Huang et al., 2021). Encouraged by this observation, we did single-cell RNA-seq for rCREG treated shrimp hemocytes to further explore its function. To collect shrimp circulating hemocytes as complete as possible, we applied iodixanol gradient centrifugation to concentrate the hemocytes for the Gel Bead-In-Emulsions (GEMs) preparation(Tattikota et al., 2020)(*Figure 1A*). A total 34693 cells including control (12544), rEGFP-treated (12640) and rCREG-treated (9509) were retained for further analyses. The 12544 control cells, 12640 rEGFP-treated cells and 9509 rCREG-treated cells exhibited a median of 5656, 6837, 7916 transcripts and 1089.5, 1245, 1364 genes per cell separately (*Figure supplement 1*). The rCREG-treated samples had the highest UMIs (unique molecular identifiers) and detected genes compared with another two groups. This observation consisted with our previous conclusion that CREG was a hemocyte activation factor.

**Figure 1.**
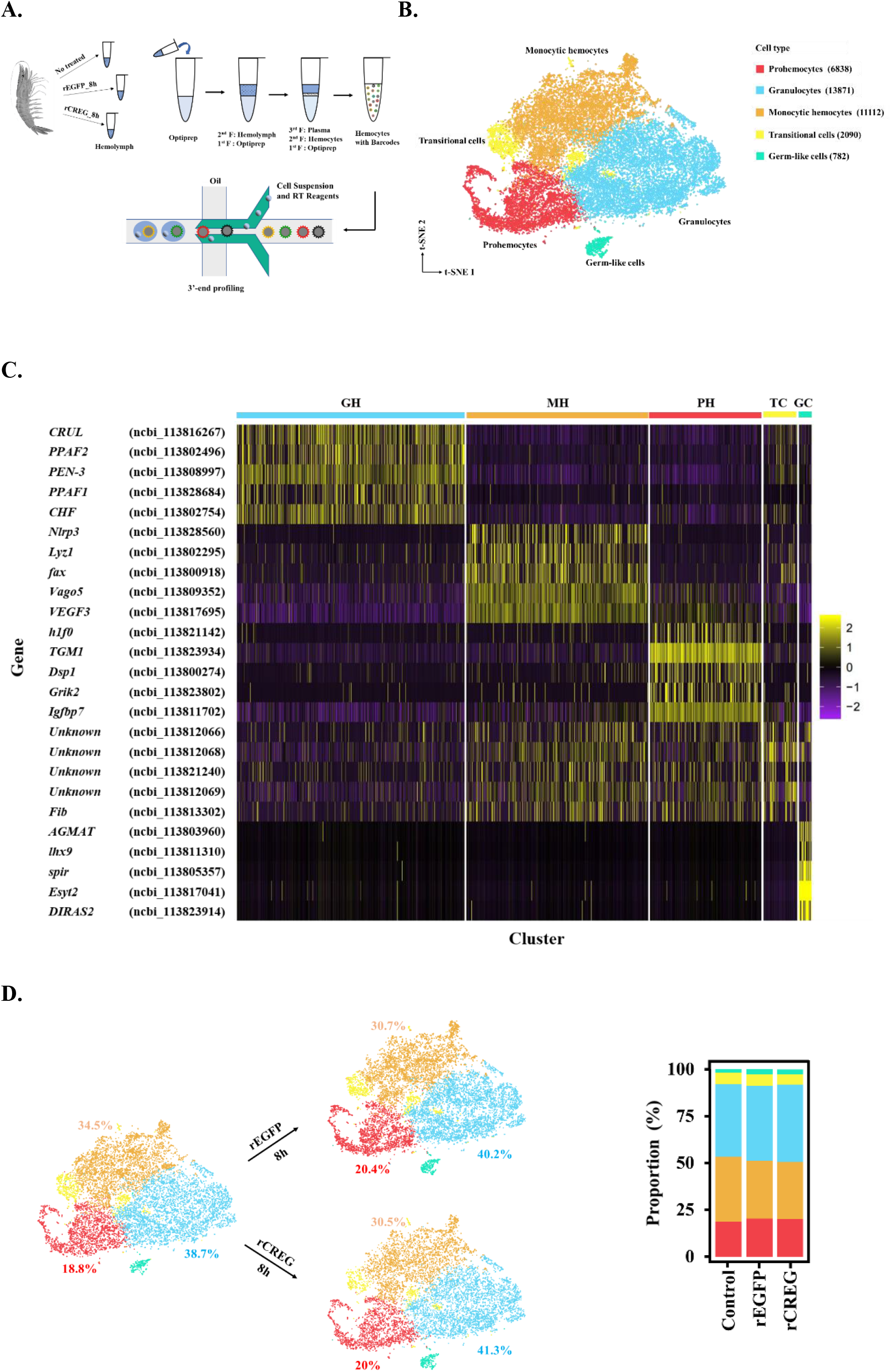
Major cell types identified in shrimp hemolymph. (**A**) A schematic workflow of sample preparation. The hemocytes were collected from non-treated, rEGFP treated and rCREG treated shrimps separately (n=15 for each treatment) and subjected for iodixanol gradient centrifugation and ScRNA-seq using Drop-seq. (**B**) A t-SNE plot showing five major cell types identified in scRNA-seq dataset (n=34693 in total; Control, 12544; rEGFP treated, 12640; rCREG treated, 9509 cells). The count of each cell type is indicated in parentheses. (**C**) A heatmap showing five representative marker genes for each major cluster. The gene name and its NCBI GeneID were listed (left) and its expression level in each cell was shown with different colors (right). (**D**) Two-dimensional projections, and proportions of the cell types for each treatment. Proportions of prohemocytes (red), monocytic hemocytes (brown) and granulocytes (blue) are indicated (left). Proportions of all five major cell types in each treatment are indicated (right).

To further define the major cell types of shrimp circulating hemocytes, we combined all 34693 cells from different treatments and applied the Canonical Correlation Analysis(Stuart et al., 2019) to do the batch correction. After that, we aggregated cell clusters and identified five major groups of isolated hemocytes including prohemocytes (PHs) (6838, 19.7%), granulocytes (GHs) (13871, 40%), monocytic hemocytes (MHs) (11112, 32%), transitional cells (TCs) (2090, 6%) and germ-like cells (GCs) (782, 2.3%), which were annotated according to their potential functions implied by the marker genes (*Figure 1B*). For granulocyte, it is featured with its ProPO system(M. Sun, Li, Zhang, Xiang, & Li, 2020). Here we identified that prophenoloxidase activating factor 1 and 2 (*PPAF1* and *PPAF2*) were highly expressed in this population. Besides ProPO system genes, secreted proteins including crustin-like protein (*CRUL*), penaeidin 3a.1 (*PEN-3*) and crustacean hematopoietic factor-like protein (*CHF*) were also highly expressed in this population (*Figure 1C; figure supplement 2A*). For monocytic hemocyte, we named this population as monocytic hemocyte because this group of hemocytes shared some critical genes with mammalian monocyte. For example, NOD-like receptor protein 3 (*Nlrp3*) is the key component for inflammasome and is highly expressed in monocyte/macrophage for processing of IL1ß(He, Hara, & Nunez, 2016). Lysosome (*Lyz1*), another canonical anti-bacterial enzyme, is a well-known macrophage secreting hydrolase(Short, Nickel, Schmitz, & Renkawitz, 1996). *Vago5*, an IFN-like anti-viral cytokine, plays its roles during anti-WSSV process(C. Li, Yang, Hong, Zhao, & Wang, 2021)(*Figure 1C; figure supplement 2B*). In general, this group of cells were featured with their secretion of anti-bacterial and anti-viral effectors. For prohemocyte, histone1 (*h1f0*) is the key component for heterochromatin assembly and its high expression is associated with less gene expression and cell stemness properties(Pan & Fan, 2016). Hemocyte transglutaminase (*TGM1*) has been identified as an immature hemocyte marker in previous study(Koiwai et al., 2021). *Igfbp7* has been shown to promote hemocyte proliferation in small abalone Haliotis diversicolor(Wang et al., 2015)(*Figure 1C; figure supplement 2C*). For transitional cell, we found that it was difficult to characterize this group of cells due to lack of exclusively expressed genes. We listed the top 5 significantly upregulated genes and showed top 3 significantly upregulated genes’ distribution (*Figure 1C; figure supplement 2D*), which indicated that this group of cells had no significant marker gene. Thus, we called this group of cells as transitional cells. For germ-like cell, we found some reproduction-related genes in this group of cells. For example, *lhx9* has been shown as a key transcription factor in gonadal development(Balasubramanian, Bui, Xie, Deng, & Gan, 2014). *spir* localization is critical for mouse oocyte asymmetric division(Jo et al., 2019)(*Figure 1C; figure supplement 2E*). Hence, we annotated this group of cells as germ-like cells.

To further answer whether CREG was a differentiation factor for shrimp hemocytes, we examined the ratio of five annotated major cell types in different treatments. The recombinant protein injection could slightly increase the proportion of granulocytes and prohemocytes and decrease the ratio of monocytic hemocytes (*Figure 1D*). However, there were no significant differences between rEGFP treatment and rCREG treatment, which suggest that CREG was probably an activation factor instead of a differentiation factor for shrimp hemocytes.

### Subtyping of shrimp immune cell clusters and construction of their differentiation trajectory

Because we are interested to figure out the immune cell classification in shrimp hemolymph, the “Transitional cell” that is lack of markers and “Germ-like cell” that is not a typical immune cell will not be further explored in the following study (*Tables supplement 1-2*). To further trace the immune cell lineages in shrimp hemolymph, we subtyped the three major classes of cells including prohemocyte, monocytic hemocyte and granulocyte, and each major type can be divided into two subtypes and these subtypes were labelled as PH1(1577, 4.5%), PH2(5261, 15.2%), MH1(10463, 30.2%), MH2(649, 1.9%), GH1(10353, 29.8%) and GH2(3518, 10.1%) separately (*Figure 2A*). Here we found a lot of unique marker genes for those subpopulations (PH1, GH2, MH2) located at the edge of t-SNE map (*Figure 2A;* *figure supplement* 3 *and tables supplement 3-5*), but cannot identify exclusive marker genes for those subpopulations (PH2, GH1, MH1) constituting of the main body of t-SNE map, the marker genes for PH2, GH1, MH1 were also highly expressed in PH1, GH2, MH2 separately (*Figure 2B; tables supplement 6-8*). Thus, we considered that these six subtypes of hemocytes probably had lineage differentiation relationships among each other. To explore this, we did the cell cycle analyses for these six subtypes of hemocyte. PH1 was featured with high expression of all marker genes for G1, G2 and M stage (*Figure 2C*). This observation was consistent with previous report that around 2-5% of circulating hemocytes were proliferating hemocytes which could be labelled with BrdU(R. Sun et al., 2013). Thus, we set PH1 as the initiating cells and applied monocle to construct differentiation trajectory for PH, GH and MH. Two major branches including monocytic hemocyte lineage and granulocyte lineage were identified to be differentiated from one common prohemocyte (*Figure 2D-E*). This observation shares common features with human myeloid cell development in which granulocyte-monocyte progenitor (GMP) differentiates into monocyte and granulocyte(Bassler et al., 2019). To further compare myeloid differentiation between shrimp and human, we screened the shrimp homologs of human myeloid differentiation related transcription factors (TFs) because TFs are key regulators for cell fate determination(Friedman, 2002). Totally 3790 differentially expressed genes among different branches were identified and shown as a specific heatmap, on which three shrimp homologs of human TFs were labelled (*Figure 2F*). *Fli1* is specifically expressed in granulocyte lineage, which is consistent with previous observation that *Fli1* deletion decreases granulocytic cell number in mice (Starck et al., 2010). *MAF* and *c-Rel* are highly expressed in monocytic hemocyte lineage (*Figure 2G*). *c-Rel* is a key TF in NF-κB pathway, which has been identified to play important roles in monocyte differentiation*(T. Li et al., 2020). MAF* is a bZip TF that could induce monocytic differentiation*(Kelly, Englmeier, Lafon, Sieweke, & Graf, 2000)*. In general, our data indicated that some myeloid regulators were conserved between shrimp and human.

**Figure 2.**
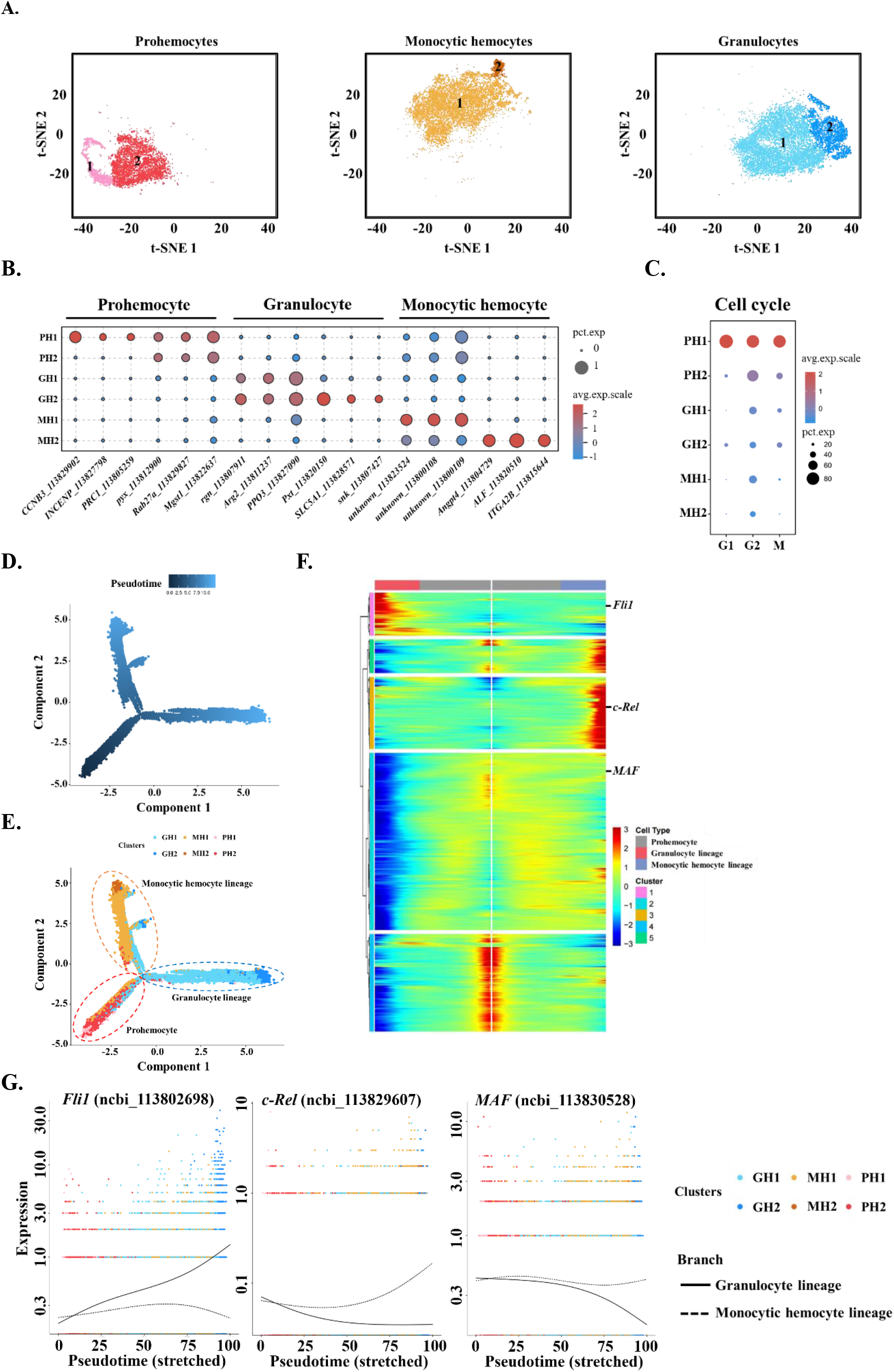
Subclustering and pseudotime trajectory analyses of three major hemocyte types in shrimp hemolymph. (**A**) Subclusters of hemocytes-prohemocytes, monocytic hemocytes, granulocytes-are projected onto two-dimensional t-SNE plots. The numbers in the plots represent the subcluster number. (**B**) Dot plot showing corresponding expression of cluster marker genes. The color indicates mean expression and dot size represents the percentage of cells within the cluster expressing the marker. Last nine digits of each marker gene are the NCBI GeneID. (**C**) Expression of cell-cycle regulating genes in 6 subtypes. Dot color indicates average expression levels and dot size displays the average percentage of cells with cell cycle controlling genes (*Cdk1*(ncbi_113818305), *CycD*(ncbi_113814652) and *CycE*(ncbi_113822658) for G1; *stg*(ncbi_113800052), *CycA*(ncbi_113821735) and *CycB*(ncbi_113803283) for G2; *polo*(ncbi_113805901), *aurB*(ncbi_113827838) and *birc5*(ncbi_113828653) for M) in each subcluster. (**D**) A differentiation trajectory of PH, GH and MH subpopulation using Monocle2 (n=31821). (**E**) Differentiation trajectory reconstruction with 6 subclusters. Prohemocyte lineage, granulocyte lineage and monocytic hemocyte lineage were labelled with red, blue, and brown circles respectively. (**F**) A heatmap showing differentially expressed gene dynamics during hemocyte differentiation process. (**G**) Spline plots showing the expression dynamics of *Fli1, c-Rel* and *MAF*. Imaginary line, monocytic hemocyte lineage; Full line, granulocyte lineage.

### Identification of a macrophage-like phagocytic cell population in shrimp hemolymph

Next, we further asked whether MH2 was a terminal differentiated monocyte like macrophage or dendric cell. Recently, the Human Cell Atlas has mapped most of gene expression across major human cells(Karlsson et al., 2021). We compared MH2 marker genes with the human database and found that human homologs of nine MH2 marker genes including chitotriosidase (*CHIT1*), lysozyme (*Lyz1)*, lipase (*LIPF*), legumain (*LGMN*), *Nlrp3*, alpha-N-acetylgalactosaminidase (*NAGA*), zinc finger E-box-binding homeobox 1(*zfh1*), caspase1 (*Casp1*) and NPC intracellular cholesterol transporter 2 (*NPC2*) were also specifically expressed in human macrophage (*Figure 3A; figures supplement 2B and 4*)(Karlsson et al., 2021), which strongly suggest that MH2 was probably the invertebrate homolog of human macrophage. To further prove this hypothesis, we labelled phagocytic hemocytes via injection of fluorescein isothiocyanate-conjugated *Vibrio Parahemolyticus* (FITC-VP), the hemocytes which engulfed FITC-VP were isolated by a cell sorter and labelled as R1 (fluorescence intensity>2×10^3^), the hemocytes with low fluorescence (<10^3^) were labelled as R2 (*Figure 3B*). To characterize what these phagocytic cells (R1) were, we quantified *CHIT1, Lyz1* and *NAGA* expression in R1 and R2 by qPCR, which indicated that these three genes expressed higher in phagocytic hemocytes (R1) than control hemocytes (R2) (*Figure 3C; table supplement 9*). In addition, we checked LYZ1, NAGA and NLRP3 by immunoblot and found that these three proteins expressed significantly higher in phagocytic hemocytes (R1) than control hemocytes (R2) (*Figure 3D*). Thus, our results indicated that the phagocytic cells in shrimp hemolymph specifically expressed MH2 marker genes. In general, our data suggest that MH2 is an invertebrate homolog of human macrophage.

**Figure 3.**
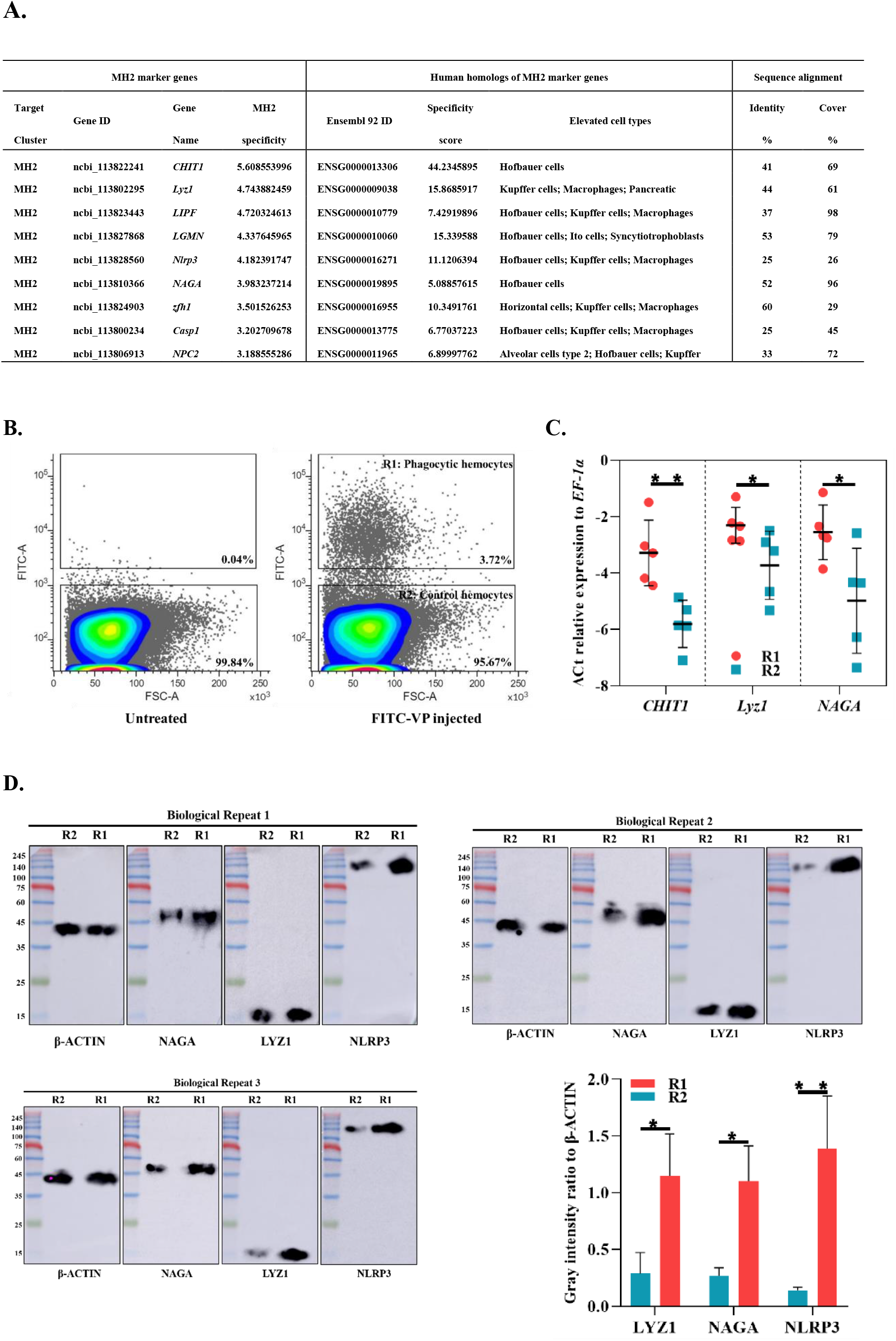
Identification of MH2 as macrophage-like phagocytic cells. (**A**) Comparison between MH2 and human macrophage marker genes. (**B**) A representative contour plot of shrimp hemocytes against FITC-VP. Threshold intensity (FITC-A) was set to <10^3^ representing control hemocytes (R2), and >2×10^3^ representing phagocytic hemocytes (R1). Phagocytic hemocytes (R1) and control hemocytes (R2) were sorted based on the forward scatter (FSC) and fluorescence intensity (FITC) two-dimensional space. (**C**) Differential gene expression analysis (*CHIT1, Lyz1* and *NAGA*) between phagocytic hemocytes (R1) and control hemocytes (R2) sorted by FACS and analyzed by qPCR. (**D**) Differential protein expression analysis (NAGA, LYZ1 and NLRP3) between phagocytic hemocytes (R1) and control hemocytes (R2) sorted by FACS. The immunoblot signals were quantified with ImageJ. The relative immunoblot signal intensities of NAGA, LYZ1 and NLRP3 compared with ß-actin were recorded with bar chart. Both qPCR and immunoblot data were analyzed by the Student *t* test and the *p* values shown in the figures are represented by \**p*<0.05 and \*\**p*<0.01.

### Comparison between hyalinocyte, semi-granulocyte, granulocyte, and the classification in this study

Next, we tried to compare our classification with traditional classification. Previous shrimp hemocytes have been divided into three major types: hyalinocyte, semi-granulocyte and granulocyte based on morphological criteria and functional properties(Soderhall, 2016). More recently, these three major types were separated with cell sorting or Percoll density gradient centrifugation and their marker genes were identified and validated by qPCR(M. Sun et al., 2020; Yang, Lu, Chen, Liao, & Chen, 2015). Here we analyzed the distribution of previous published marker genes including lysosome membrane protein2 (*LIMP2*, ncbi_113826216), tubulin beta chain (*TUBB4B*, ncbi_113826677), dipeptidyl peptidase 1 (*CTSC*, ncbi_113824311), transglutaminase 1 (*TGM1*, ncbi_113823934) for hyalinocyte (*Figure 4A; figure supplement 2C*); beta-arrestin-1 (*ARRB1*, ncbi_113804686), ADP-ribosylation factor 6 (*ARF6*, ncbi_113820333), lysozyme (*Lyz1*, ncbi_113802295), Penaeid-3a (*PEN-3*, ncbi_113808997) for semi-granulocyte (*Figure 4B; figure supplement 2A-B*); clone ZAP 18 putative antimicrobial peptide (*CRU*, ncbi_113801825), phenoloxidase-activating factor 3 (*PPAF3*, ncbi_113800184), phenoloxidase 3-like (*PPO3*, ncbi_113827090), peroxinectin (*Pxt*, ncbi_113820150) for granulocyte (*Figure 4C; figure supplement 3C*). The hyalinocyte marker genes were highly expressed in PH1, PH2, MH1 and MH2, which included prohemocytes and monocytic hemocytes (*Figure 4A and D*). The semi-granulocyte marker genes were highly expressed in GH2 and MH2, which included monocytic hemocytes and granulocytes (*Figure* 4*B* *and D*). These data were consistent with previous observation that hyalinocytes contained both proliferating progenitors and phagocytic cells(Soderhall, 2016) and explained that why some studies also observed that semi-granulocytes had phagocytic activities(M. Sun et al., 2020). The granulocyte marker genes were consistent with our observation and highly expressed in GH2 (*Figure 4C-D*), which indicated that the granulocyte was indeed the largest cell type with condensed granules inside(Soderhall, 2016).

**Figure 4.**
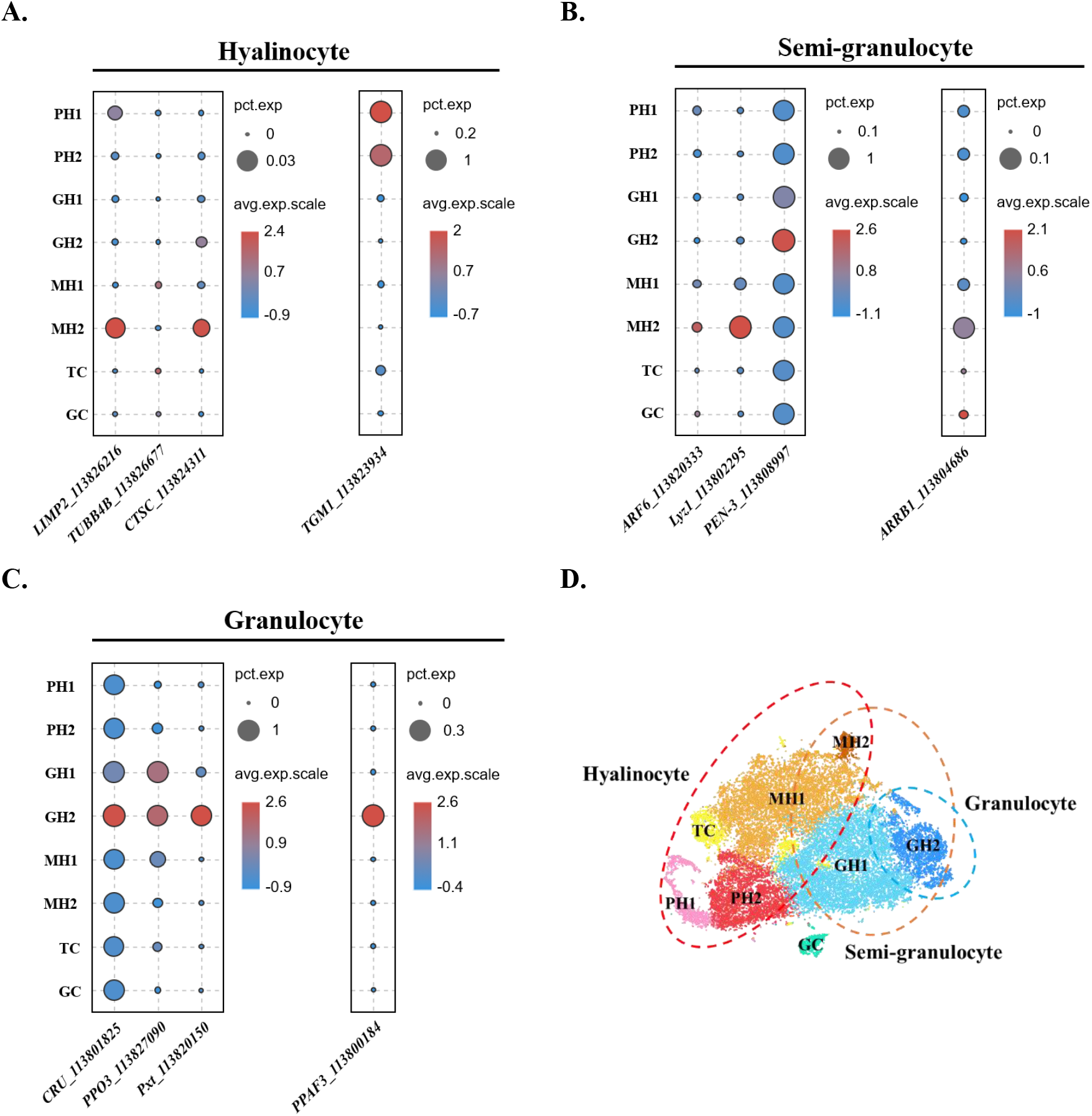
Comparison between the traditional classification and the classification in this study. **A**) Dot plot showing corresponding expression of previously reported hyalinocyte marker genes in eight subclusters. (**B**) Dot plot showing corresponding expression of previously reported semi-granulocyte marker genes in eight subclusters. (**C**) Dot plot showing corresponding expression of previously reported granulocyte marker genes in eight subclusters. The color indicates mean expression and dot size represents the percentage of cells within the cluster expressing the marker. (**D**) A proposed model for comparison between two classifications. The hyalinocyte, semi-granulocyte and granulocyte were labelled on the t-SNE map with red, brown, and blue circles respectively.

## Discussion

Myeloid cells play important roles for animal to adapt complex and volatile environment. Their fast responsive ability protects animal from various pathogen invasion and shapes host-microbiota symbiosis(Bassler et al., 2019). To maintain this symbiosis, coevolution between myeloid cells-microbiota lasts throughout the whole metazoan evolutionary history, which suggest that some common ways to restrict microbes are highly conserved. Here our data reveal a possibility that phagocytic cell and ROS generating cell are probably two major immune cell lineages co-evoluting with metazoan to effectively maintain host-microbiota balance. In addition, *Nlrp3* and *Casp1* were identified in shrimp phagocytic cells, which strongly suggest that inflammasome-mediated pyroptosis is probably a common strategy developed by metazoan for quickly defensing invaded pathogens(Wang et al., 2021).

Phagocytic ability is one of the most fundamental function for organism. Unicellular organism maintains its life via phagocytosis. The cells in multicellular organism have functional specializations, which increase life adaptability. Thus, phagocytic ability in metazoan is limited in certain cells. The ratio of phagocytic cells in different species varies a lot. For example, around 40% to 60% hemocyte could engulf pathogens in primitive oyster *Crassostrea gigas*(J. Sun et al., 2021). While in fish red blood cells, major components of immune cells, possess phagocytic activity to invaded pathogens(Xu et al., 2021). In human, most of myeloid cells including monocyte, macrophage, dendritic cell, and neutrophil could engulf pathogens(Bassler et al., 2019). Crustacean seems possess less phagocytic cells compared with other species, which may be due to its unique endosymbiosis with microbes in its hemolymph, which has been partially unveiled by recent studies(Gong et al., 2021; Zheng et al., 2021)

Crustacean hemocytes have been classified into hyalinocytes, semi-granulocytes and granulocytes based on morphology and function for the past decades(Soderhall, 2016). We compared our classification with the traditional one in this study and explained some debates in this field. For example, why hyalinocytes have both proliferating activity and phagocytic activity(Soderhall, 2016). Why semi-granulocytes have phagocytic activity(M. Sun et al., 2020). In general, we classified crustacean immune cell into prohemocyte, monocytic hemocyte and granulocyte and showed its similarity with vertebrate myeloid cells, which is the promising evidence for vertebrate myeloid cell evolution from marine invertebrate.

## Materials and methods

### Experimental organisms

The shrimps were purchased from Shantou local farms. Shrimps were cultured in water tanks filled with aerated seawater at 20°C upon delivery and acclimatized for 2-3 days before the experiments. All the animal-related experiments were in accordance with Shantou University guidelines.

### Recombinant proteins preparation and injection

The recombinant EGFP and CREG were purified as previously described(Huang et al., 2021). Sixty shrimps were equally divided into three groups. One group was left without treatment and labelled as a control. Another two groups were injected with rEGFP or rCREG (1 μg/g) separately. The hemolymph was collected at 8 hours post injection and mixed well for each group. 1.5 mL hemolymph was loaded onto OptiPrep (Axis-shield, NO) separation solution (1.09 g/mL) and centrifuged at 2000 rpm for 10 min at 4°C. The circulating hemocytes were concentrated between hemolymph and separation solution and carefully collected with a pipettor. The collected hemocytes were stained with 0.4% Trypan blue to estimate the cell viability. After that, the cells with>85% viability were subjected for further experiment.

### Preparation of single-cell cDNA library

The hemocyte suspensions were loaded on a 10X Genomics GemCode Single-cell instrument to generate single-cell gel bead-in-emulsions (GEMs). Libraries were generated with Chromium Next GEM Single Cell 3’Reagent Kits v3.1 (10X Genomics, USA). In brief, the hemocytes in GEMs were mixed with primers including an Illumina® R1 sequence, barcodes, Unique Molecular Identifiers (UMIs) and poly-dT primers. The full-length cDNA libraries were then reverse transcribed from each GEM and constructed via End Repair, A-tailing, Adaptor Ligation and PCR.

### ScRNA sequencing

The constructed libraries were sequenced with an Illumina Sequencer. In brief, the Single Cell 3’ 16 bp 10X Barcode and 10 bp UMI were encoded in Read 1, while Read 2 was used to sequence the cDNA fragment. Sample index sequences were incorporated as the i7 index read. Read 1 and Read 2 were standard Illumina® sequencing primer sites used in paired-end sequencing.

### Bioinformatics analyses

Raw BCL files was converted into FASTQ files by 10X Genomics Cell Ranger software (version 5.0). After that, the reads were mapped to the shrimp genome (taxid: 6689), and the reads uniquely intersecting an exon at least 50% were considered for UMI counting. The valid barcodes were identified based on the EmptyDrops method(Lun et al., 2019). The hemocytes by gene matrices for control, rEGFP treatment and rCREG treatment were individually imported to Seurat version 3.1.1 for the following analyses(Butler, Hoffman, Smibert, Papalexi, & Satija, 2018).

The cells with UMIs (≥17000), mitochondrial gene (≥10%), ≤230 detected genes or ≥2200 detected genes were filtered out. The qualified cells were normalized via “LogNormalize” method, which normalizes the gene expression for each cell by the total expression. The formula is shown as follows:

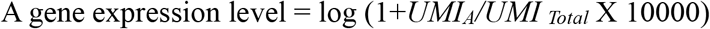

The batch effect was corrected with the Canonical Correlation Analysis(Stuart et al., 2019). Integrated expression matrix was then scaled and performed on principal component analysis (PCA) for dimensional reduction. After that, the significant principal components (PCs) were identified as those who had a strong enrichment of low p-value genes(Chung & Storey, 2015).

### Phagocytic cell labelling and sorting

Shrimp phagocytic cells were labelled as previously described(Huang et al., 2021). In brief, FITC labelled *Vibrio parahaemolyticus* (VP) (2×10^6^ particles/g) were injected into shrimps. The hemocytes were collected from thirty to forty shrimps at 2 hours post injection for each sorting which was performed on a FACSMelody cell sorter (BD Biosciences, USA). The fluorescence boundary was set based on detection of the shrimp hemocyte self-fluorescence without VP injection.

### Sorted hemocyte RNA and protein collection for RT-qPCR and immunoblot analyses

For each experiment, 50k to 100k events from the phagocytic hemocytes (R1) and the control hemocytes (R2) were collected respectively. Total RNA from collected samples was purified with the RNAprep Pure Micro Kit (TIANGEN, Beijing, China) and reversely transcribed into cDNA with a First Strand cDNA Synthesis Kit (Beyotime, Shanghai, China). QPCR was performed as previously described(Luo et al., 2022)(*Table supplement* 10), the gene expression level was recorded as relative expression to *EF-1α*. This experiment was biologically repeated for five times. Total proteins from sorted hemocyte were precipitated by adding with 1/100 volume of 2% Na deoxycholate (Macklin, Shanghai, China) and 1/10 volume of 100% trichloroacetic acid (Macklin, Shanghai, China), followed by vortex and centrifugation at 15000 g for 15 min at 4°C. The pellet was collected for the following SDS-PAGE and immunoblot as described before(Luo et al., 2022). This experiment was biologically repeated for three times. The antibodies were obtained: ß-ACTIN (AF5003, Beyotime, Shanghai, China), anti-NAGA (13686-T24, SinoBiological, Beijing, China), anti-LYZ1 (bs-0816R, Bioss Antibodies, MA, USA). The polypeptide antibody against shrimp NLRP3 (aa29-42) was prepared by GenScript (Nanjing, China).

## Acknowledgments

This work was supported by National Natural Science Foundation of China (41976123 to F.W); the Sail Plan Program for the Introduction of Outstanding Talents of Guangdong Province of China (14600703 to F.W); 2020 Li Ka Shing Foundation Cross-Disciplinary Research Grant (2020LKSFG01E to Y.Z).

## Data Availability

The sequence data reported in this paper have been deposited in the genome sequence archive of Beijing Institute of Genomics, Chinese Academy of Sciences, gsa.big.ac.cn (accession no. PRJCA006297). And all other data have been made available in this manuscript and in the Supplementary Material online.

## Supplementary Figures

**Figure supplement 1.**
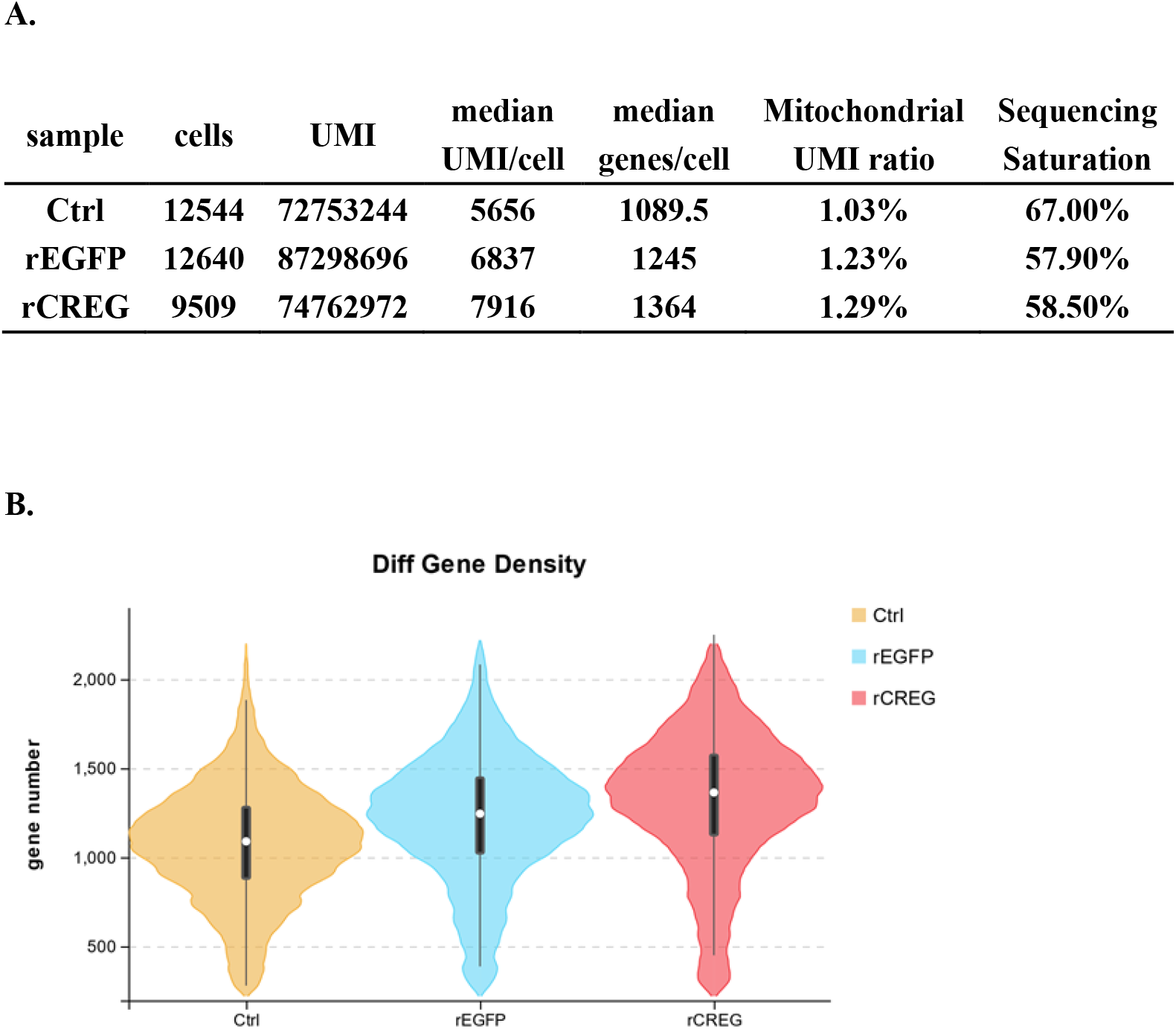
Overall quality of single-cell transcriptomic data.

**Figure supplement 2.**
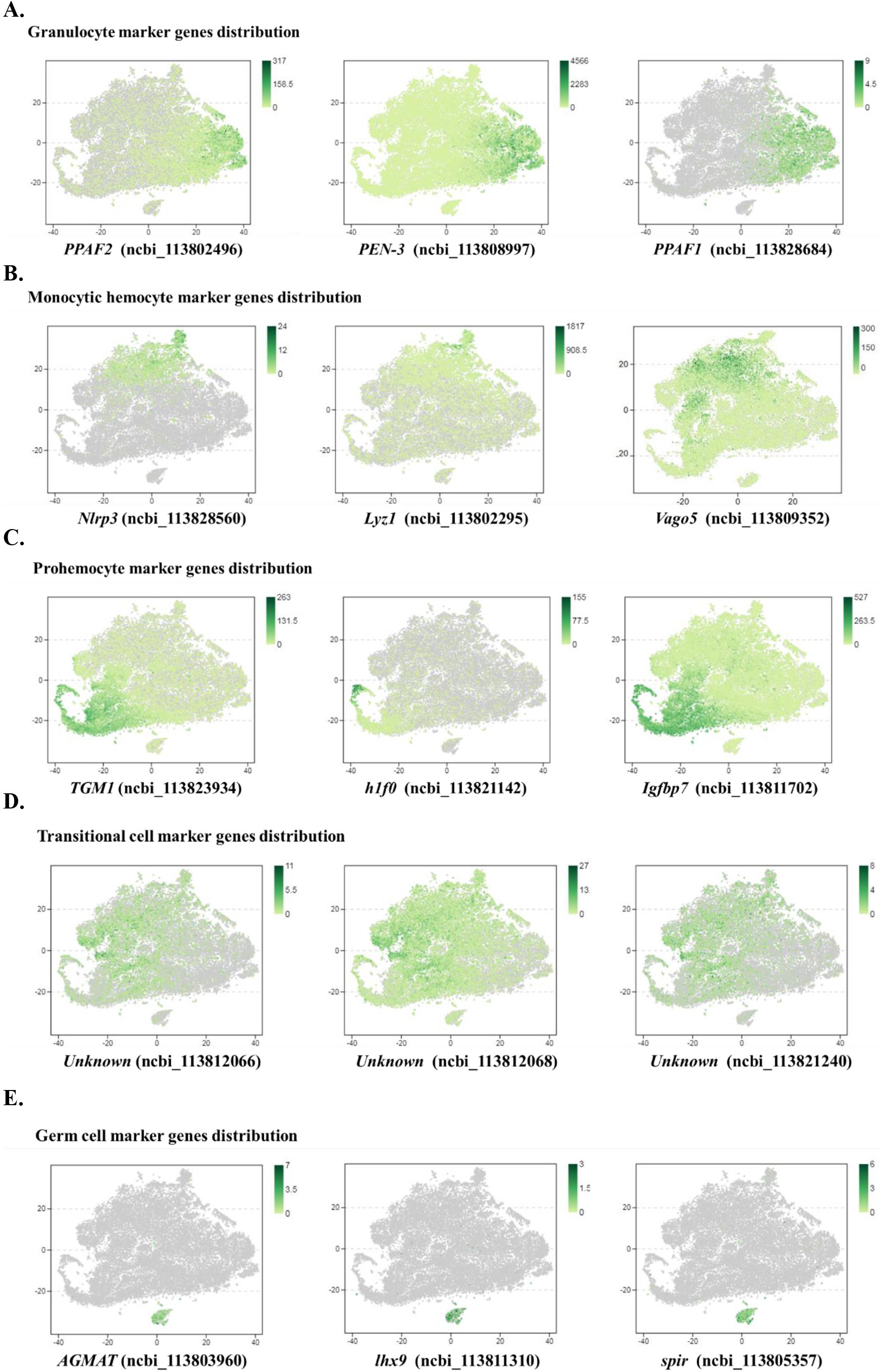
Distribution of the marker genes from major cell types.

**Figure supplement 3.**
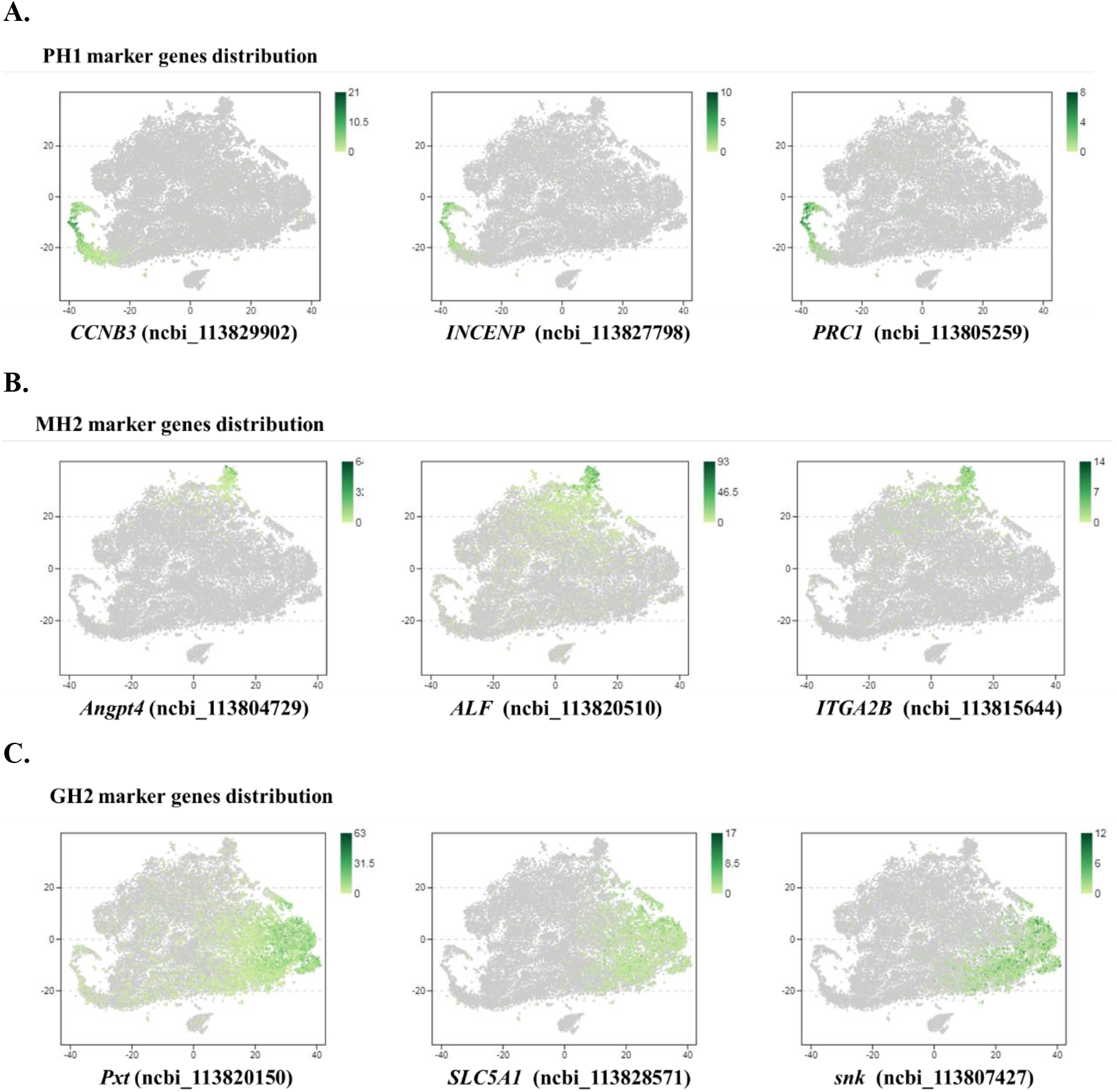
Distribution of the marker genes for PH1, MH2 and GH2.

**Figure supplement 4.**
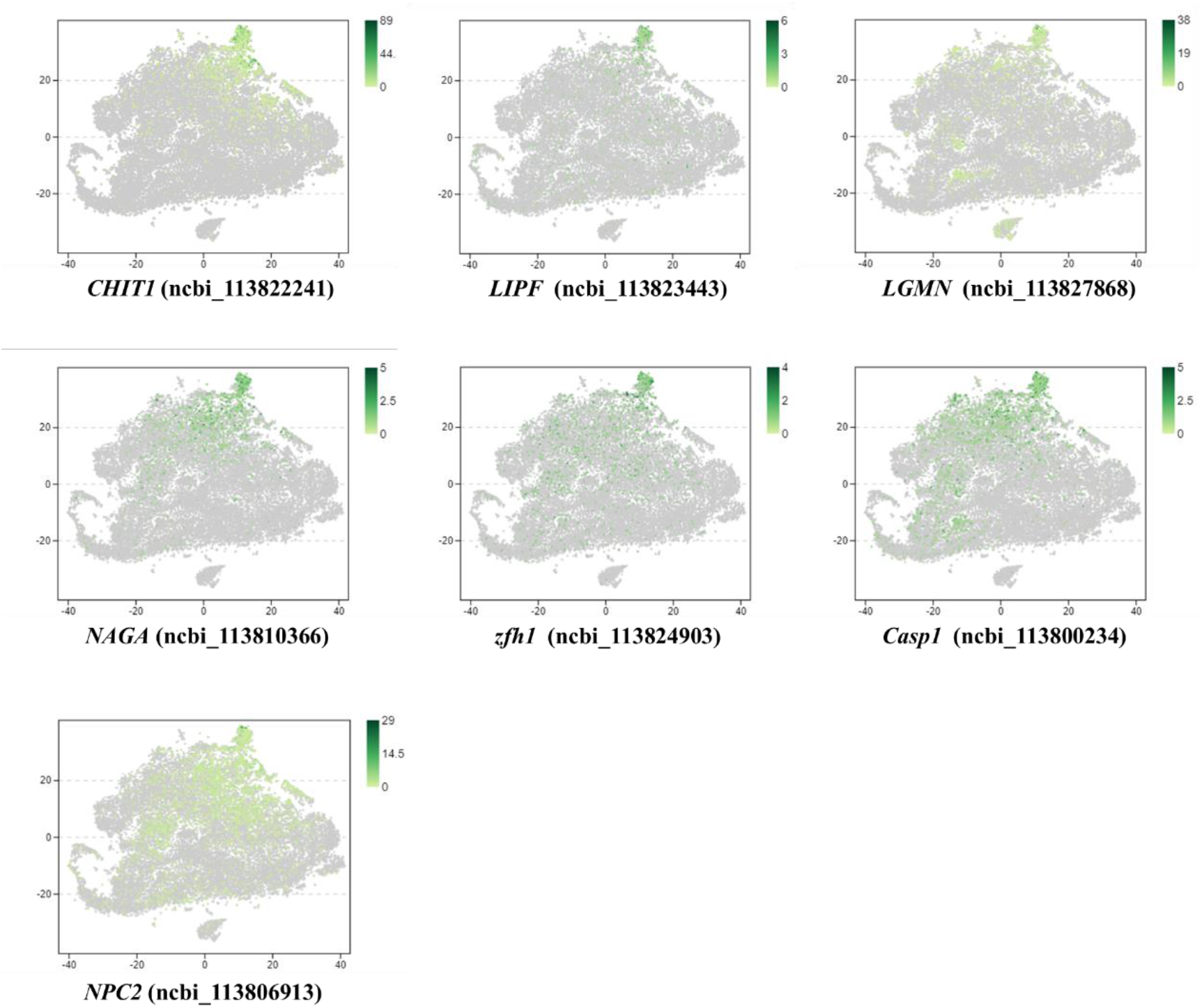
Distribution of MH2 marker genes which are conserved with that of human macrophages.

## Supplementary Tables

**Table supplement 1.**
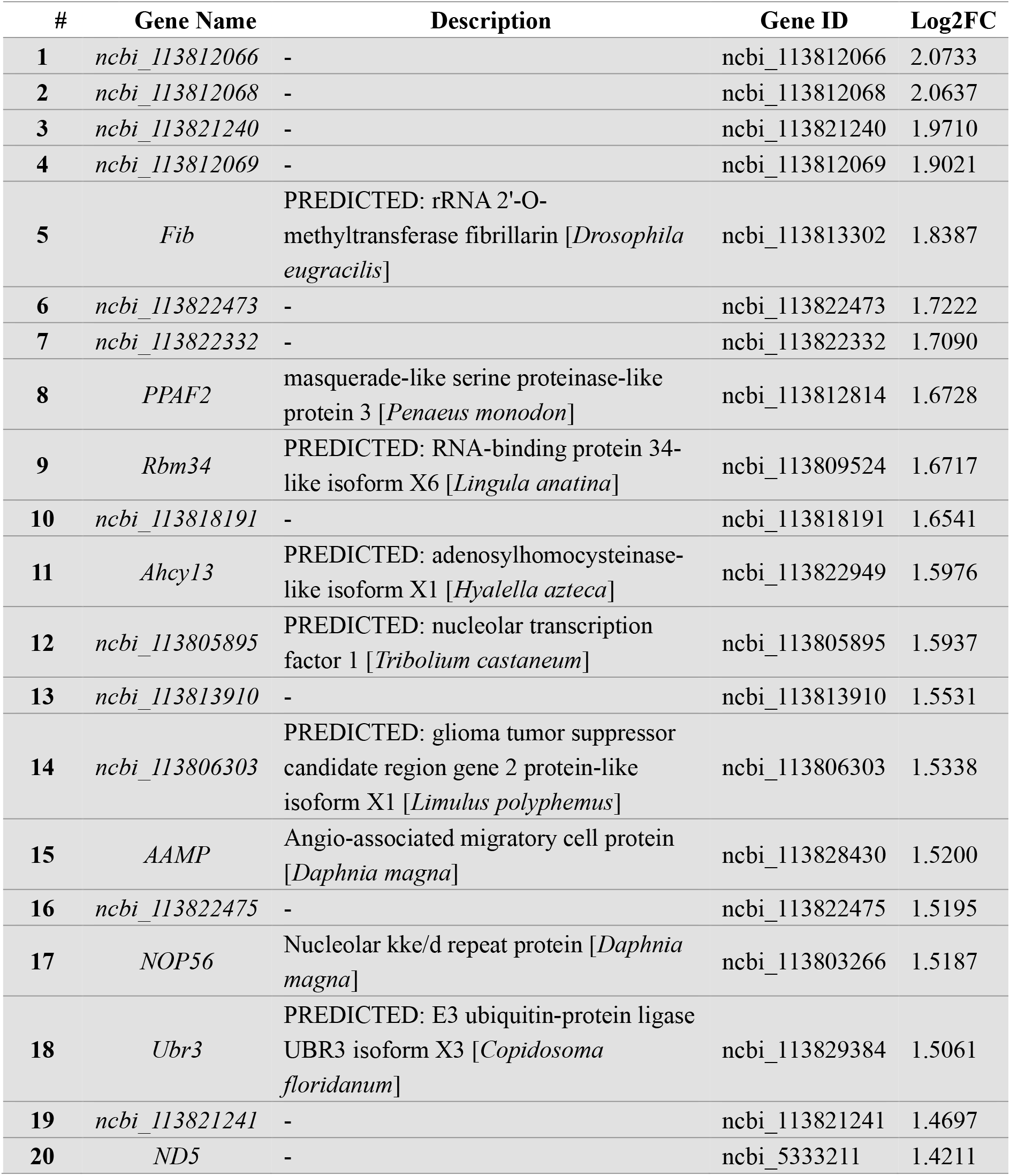
Top 20 specifically expressed marker genes for transitional cell (TC).

| # | Gene Name | Description | Gene ID | Log2FC |
| --- | --- | --- | --- | --- |
| 1 | *ncbi_113812066* | - | ncbi_113812066 | 2.0733 |
| 2 | *ncbi_113812068* | - | ncbi_113812068 | 2.0637 |
| 3 | *ncbi_113821240* | - | ncbi_113821240 | 1.9710 |
| 4 | *ncbi_113812069* | - | ncbi_113812069 | 1.9021 |
| 5 | *Fib* | PREDICTED: rRNA 2'-O-methyltransferase fibrillarin [*Drosophila eugracilis*] | ncbi_113813302 | 1.8387 |
| 6 | *ncbi_113822473* | - | ncbi_113822473 | 1.7222 |
| 7 | *ncbi_113822332* | - | ncbi_113822332 | 1.7090 |
| 8 | *PPAF2* | masquerade-like serine proteinase-like protein 3 [*Penaeus monodon*] | ncbi_113812814 | 1.6728 |
| 9 | *Rbm34* | PREDICTED: RNA-binding protein 34-like isoform X6 [*Lingula anatina*] | ncbi_113809524 | 1.6717 |
| 10 | *ncbi_113818191* | - | ncbi_113818191 | 1.6541 |
| 11 | *Ahcy13* | PREDICTED: adenosylhomocysteinase-like isoform X1 [*Hyalella azteca*] | ncbi_113822949 | 1.5976 |
| 12 | *ncbi_113805895* | PREDICTED: nucleolar transcription factor 1 [*Tribolium castaneum*] | ncbi_113805895 | 1.5937 |
| 13 | *ncbi_113813910* | - | ncbi_113813910 | 1.5531 |
| 14 | *ncbi_113806303* | PREDICTED: glioma tumor suppressor candidate region gene 2 protein-like isoform X1 [*Limulus polyphemus*] | ncbi_113806303 | 1.5338 |
| 15 | *AAMP* | Angio-associated migratory cell protein [*Daphnia magna*] | ncbi_113828430 | 1.5200 |
| 16 | *ncbi_113822475* | - | ncbi_113822475 | 1.5195 |
| 17 | *NOP56* | Nucleolar kke/d repeat protein [*Daphnia magna*] | ncbi_113803266 | 1.5187 |
| 18 | *Ubr3* | PREDICTED: E3 ubiquitin-protein ligase UBR3 isoform X3 [*Copidosoma floridanum*] | ncbi_113829384 | 1.5061 |
| 19 | *ncbi_113821241* | - | ncbi_113821241 | 1.4697 |
| 20 | *ND5* | - | ncbi_5333211 | 1.4211 |

**Table supplement 2.** Top 20 specifically expressed marker genes for germ-like cell (GC).

| # | Gene Name | Description | Gene ID | Log2FC |
| --- | --- | --- | --- | --- |
| 1 | *ncbi_113826700* | - | *ncbi_113826700* | 12.6604 |
| 2 | *ncbi_113813962* | homeobox domain protein [*Dictyocaulus viviparus*] | *ncbi_113813962* | 12.0217 |
| 3 | *ncbi_113830473* | - | *ncbi_113830473* | 11.9115 |
| 4 | *ncbi_113816939* | - | *ncbi_113816939* | 11.8305 |
| 5 | *ncbi_113813667* | - | *ncbi_113813667* | 11.5591 |
| 6 | *ncbi_113823051* | - | *ncbi_113823051* | 11.4330 |
| 7 | *ncbi_113805945* | - | *ncbi_113805945* | 11.3896 |
| 8 | *ncbi_113803297* | - | *ncbi_113803297* | 11.3623 |
| 9 | *ncbi_113802931* | PREDICTED: zinc finger protein 629-like [*Hyaella azteca*] | *ncbi_113802931* | 11.1494 |
| 10 | *AGMAT* | Agmatinase, mitochondrial [*Crassostrea gigas*] | *ncbi_113803960* | 11.1002 |
| 11 | *lhx9* | PREDICTED: LIM/homeobox protein Lhx2-like [*Thamnophis sirtalis*] | *ncbi_113811310* | 11.0221 |
| 12 | *spir* | Protein spire [*Zootermopsis nevadensis*] | *ncbi_113805357* | 10.8140 |
| 13 | *Esy2* | PREDICTED: LOW QUALITY PROTEIN: extended synaptotagmin-1-like [*Hyaella azteca*] | *ncbi_113817041* | 10.7314 |
| 14 | *DIRAS2* | PREDICTED: GTP-binding protein Di-Ras2 [*Cavia porcellus*] | *ncbi_113823914* | 10.5484 |
| 15 | *ncbi_113818891* | - | *ncbi_113818891* | 10.5066 |
| 16 | *ncbi_113807122* | - | *ncbi_113807122* | 10.5000 |
| 17 | *ncbi_113803272* | - | *ncbi_113803272* | 10.3566 |
| 18 | *aop* | PREDICTED: ets DNA-binding protein pokkuri-like [*Hyaella azteca*] | *ncbi_113824567* | 10.1863 |
| 19 | *Ptprb* | PREDICTED: receptor-type tyrosine-protein phosphatase O [*Crassostrea gigas*] | *ncbi_113821981* | 9.9991 |
| 20 | *ncbi_113820066* | - | *ncbi_113820066* | 9.8777 |

**Table supplement 3.** Top 20 specifically expressed marker genes for prohemocyte 1 (PH1).

| # | Gene Name | Description | Gene ID | Log2FC |
| --- | --- | --- | --- | --- |
| 1 | *CCNB3* | PREDICTED: G2/mitotic-specific cyclin-B3-like isoform X2 [*Hyalella azteca*] | ncbi_113829902 | 7.8924 |
| 2 | *INCENP* | PREDICTED: inner centromere protein A-like [*Lingula anatina*] | ncbi_113827798 | 7.7589 |
| 3 | *PRC1* | PREDICTED: protein regulator of cytokinesis 1-like isoform X1 [*Lingula anatina*] | ncbi_113805259 | 7.4668 |
| 4 | *ncbi_113810803* | - | ncbi_113810803 | 7.3779 |
| 5 | *Cenpe* | PREDICTED: centromere-associated protein E-like [*Hyalella azteca*] | ncbi_113813509 | 7.2860 |
| 6 | *ncbi_113810390* | PREDICTED: centromere protein F [*Clupea harengus*] | ncbi_113810390 | 7.2335 |
| 7 | *ASPM* | PREDICTED: abnormal spindle-like microcephaly-associated protein homolog isoform X1 [*Strongylocentrotus purpuratus*] | ncbi_113819387 | 7.2256 |
| 8 | *Klp61F* | PREDICTED: kinesin-like protein KIF11 [*Hyalella azteca*] | ncbi_113803105 | 7.2130 |
| 9 | *hlf0* | Histone 1 [*Macrobrachium rosenbergii*] | ncbi_113821144 | 7.0063 |
| 10 | *ncbi_113809781* | - | ncbi_113809781 | 6.9067 |
| 11 | *PBK* | PREDICTED: lymphokine-activated killer T-cell-originated protein kinase-like [*Ciona intestinalis*] | ncbi_113826652 | 6.6491 |
| 12 | *hlf0* | Histone 1 [*Macrobrachium rosenbergii*] | ncbi_113821141 | 6.6180 |
| 13 | *ncbi_113807182* | RAC-GTPase activating protein 1 [*Penaeus monodon*] | ncbi_113807182 | 6.5997 |
| 14 | *ncbi_113820084* | PREDICTED: kinetochore protein Spc24-like [*Lingula anatina*] | ncbi_113820084 | 6.3148 |
| 15 | *ncbi_113823166* | PREDICTED: disks large-associated protein 5-like [*Hyalella azteca*] | ncbi_113823166 | 6.2323 |
| 16 | *Mad2l1* | Mitotic spindle assembly checkpoint protein MAD2A [*Lepeophtheirus salmonis*] | ncbi_113827655 | 6.2180 |
| 17 | *CIT* | Citron Rho-interacting kinase [*Zootermopsis nevadensis*] | ncbi_113808119 | 6.1719 |
| 18 | *hlf0* | Histone 1 [*Macrobrachium rosenbergii*] | ncbi_113821142 | 6.1437 |
| 19 | *ncbi_113823276* | PREDICTED: chromatin assembly factor 1 subunit A-like [*Saccoglossus kowalevskii*] | ncbi_113823276 | 6.1003 |
| 20 | *SPDL1* | PREDICTED: protein Spindly-B-like [*Hyalella azteca*] | ncbi_113812106 | 5.9967 |

**Table supplement 4.** Top 20 specifically expressed marker genes for granulocyte 2 (GH2).

| # | Gene Name | Description | Gene ID | Log2FC |
| --- | --- | --- | --- | --- |
| 1 | *ncbi_113801830* | crustin-like antimicrobial peptide<br>[*Fenneropenaeus chinensis*] | ncbi_113801830 | 6.2990 |
| 2 | *PPAF3* | serine protease [*Scylla paramamosain*] | ncbi_113800184 | 6.2674 |
| 3 | *ncbi_113801828* | crustin-like antimicrobial peptide<br>[*Fenneropenaeus chinensis*] | ncbi_113801828 | 6.0105 |
| 4 | *ncbi_113801827* | crustin-like antimicrobial peptide<br>[*Fenneropenaeus chinensis*] | ncbi_113801827 | 5.9763 |
| 5 | *Pxt* | peroxinectin [*Fenneropenaeus merguensis*] | ncbi_113820161 | 5.8753 |
| 6 | *ncbi_113810905* | PREDICTED: WW domain-containing<br>protein C660.06-like [*Hyalella azteca*] | ncbi_113810905 | 5.5206 |
| 7 | *ncbi_113810909* | PREDICTED: WW domain-containing<br>protein C660.06-like [*Hyalella azteca*] | ncbi_113810909 | 5.4201 |
| 8 | *ncbi_113801821* | crustin-like antimicrobial peptide<br>[*Fenneropenaeus chinensis*] | ncbi_113801821 | 5.0945 |
| 9 | *PPAF2* | masquerade-like protein [*Pacifastacus*<br>*leniusculus*] | ncbi_113804200 | 5.0241 |
| 10 | *ncbi_113811243* | - | ncbi_113811243 | 4.9710 |
| 11 | *ENPP3* | PREDICTED: venom phosphodiesterase 2-like [*Hyalella azteca*] | ncbi_113806994 | 4.8429 |
| 12 | *ncbi_113808014* | - | ncbi_113808014 | 4.6968 |
| 13 | *PPAF2* | mas [*Litopenaeus vannamei*] | ncbi_113802496 | 4.6543 |
| 14 | *ncbi_113808543* | single whey acidic protein domain-containing protein isoform 3 [*Penaeus*<br>*monodon*] | ncbi_113808543 | 4.5382 |
| 15 | *Pxt* | peroxinectin [*Fenneropenaeus chinensis*] | ncbi_113820150 | 4.3553 |
| 16 | *ncbi_113819749* | X-linked interleukin-1 receptor accessory<br>protein-like 1 [*Zootermopsis nevadensis*] | ncbi_113819749 | 4.0583 |
| 17 | *ncbi_113816771* | - | ncbi_113816771 | 3.9953 |
| 18 | *SLC5A1* | Sodium/glucose cotransporter [*Daphnia*<br>*magna*] | ncbi_113828571 | 3.8910 |
| 19 | *lcc1* | PREDICTED: L-ascorbate oxidase-like<br>[*Anoplophora glabripennis*] | ncbi_113802330 | 3.8571 |
| 20 | *snk* | clip domain serine proteinase 3 [*Portunus*<br>*trituberculatus*] | ncbi_113807427 | 3.8212 |

**Table supplement 5.** Top 60 specifically expressed marker genes for monocytic hemocyte 2 (MH2).

| # | Gene Name | Description | Gene ID | Log2FC |
| --- | --- | --- | --- | --- |
| 1 | *Angpt4* | PREDICTED: microfibril-associated glycoprotein 4-like [*Clupea harengus*] | ncbi_113804729 | 8.3431 |
| 2 | *ncbi_113826144* | PREDICTED: receptor-type tyrosine-protein phosphatase S-like, partial [*Amphimedon queenslandica*] | ncbi_113826144 | 6.6914 |
| 3 | *ncbi_113812270* | - | ncbi_113812270 | 6.4280 |
| 4 | *ncbi_113823074* | - | ncbi_113823074 | 6.3400 |
| 5 | *ALF* | antilipopolysaccharide factor isoform 4 [*Fenneropenaeus chinensis*] | ncbi_113820510 | 6.2994 |
| 6 | *ITGA2B* | integrin alpha 5 [*Fenneropenaeus chinensis*] | ncbi_113815644 | 5.9588 |
| 7 | *ncbi_113810024* | chitin binding-like protein [*Fenneropenaeus chinensis*] | ncbi_113810024 | 5.7851 |
| 8 | *CHIT1* | chitinase 5 [*Litopenaeus vannamei*] | ncbi_113822241 | 5.6086 |
| 9 | *chic* | chicadee/profilin [*Litopenaeus vannamei*] | ncbi_113812492 | 5.5349 |
| 10 | *ncbi_113811918* | PREDICTED: galactose-specific lectin nattectin-like [*Hyaella azteca*] | ncbi_113811918 | 5.3282 |
| 11 | *ncbi_113810108* | anti-lipopolysaccharide factor isoform 6 [*Penaeus monodon*] | ncbi_113810108 | 5.2874 |
| 12 | *ncbi_113813946* | chitin binding-like protein [*Fenneropenaeus chinensis*] | ncbi_113813946 | 5.1672 |
| 13 | *ncbi_113810025* | PREDICTED: beta-2 adrenergic receptor-like [*Neolamprologus brichardi*] | ncbi_113810025 | 5.1407 |
| 14 | *fax* | PREDICTED: failed axon connections-like isoform X1 [*Hyaella azteca*] | ncbi_113800918 | 4.9016 |
| 15 | *GGH* | PREDICTED: gamma-glutamyl hydrolase-like isoform X1 [*Limulus polyphemus*] | ncbi_113829328 | 4.8150 |
| 16 | *Lyz1* | lysozyme [*Litopenaeus vannamei*] | ncbi_113802295 | 4.7439 |
| 17 | *LIPF* | lipase 3 [*Culex quinquefasciatus*] | ncbi_113823443 | 4.7203 |
| 18 | *Tl* | toll2 [*Litopenaeus vannamei*] | ncbi_113824360 | 4.6130 |
| 19 | *ncbi_113806935* | PREDICTED: LOW QUALITY PROTEIN: CD109 antigen-like [*Galendromus occidentalis*] | ncbi_113806935 | 4.5596 |
| 20 | *ncbi_113812219* | C-type lectin 3, partial [*Marsupenaeus japonicus*] | ncbi_113812219 | 4.4413 |
| 21 | *ncbi_113823707* | - | ncbi_113823707 | 4.3961 |
| 22 | *ncbi_113825861* | fibrinogen-related protein 1 isoform 5 [*Litopenaeus vannamei*] | ncbi_113825861 | 4.3609 |
| 23 | *LGMN* | PREDICTED: legumain-like [*Lingula anatina*] | ncbi_113827868 | 4.3376 |
| 24 | *ncbi_113800836* | PREDICTED: sialin-like [*Bombyx mori*] | ncbi_113800836 | 4.3028 |
| 25 | *ncbi_113818495* | PREDICTED: C-type lectin domain family 4 member M-like [*Hyaella azteca*] | *ncbi_113818495* | 4.2545 |
| 26 | *Nlrp3* | PREDICTED: NLR family CARD domain-containing protein 4-like [*Lingula anatina*] | *ncbi_113828560* | 4.1824 |
| 27 | *ncbi_113826852* | PREDICTED: uncharacterized protein LOC108669804 [*Hyaella azteca*] | *ncbi_113826852* | 4.0856 |
| 28 | *ncbi_113823075* | C-type lectin 2 [*Marsupenaeus japonicus*] | *ncbi_113823075* | 4.0405 |
| 29 | *ncbi_113802813* | Adenylate cyclase [*Daphnia magna*] | *ncbi_113802813* | 4.0027 |
| 30 | *NAGA* | Alpha-N-acetylgalactosaminidase [*Harpegnathos saltator*] | *ncbi_113810366* | 3.9832 |
| 31 | *ncbi_113829941* | - | *ncbi_113829941* | 3.9437 |
| 32 | *ncbi_113804553* | C-type lectin 2 [*Marsupenaeus japonicus*] | *ncbi_113804553* | 3.9273 |
| 33 | *ncbi_113827405* | PREDICTED: thiosulfate sulfurtransferase/rhodanese-like domain-containing protein 3 isoform X1 [*Hyaella azteca*] | *ncbi_113827405* | 3.9198 |
| 34 | *ncbi_113803502* | - | *ncbi_113803502* | 3.8626 |
| 35 | *MEGF11* | PREDICTED: multiple epidermal growth factor-like domains protein 11 [*Hyaella azteca*] | *ncbi_113823866* | 3.8543 |
| 36 | *ncbi_113817215* | PREDICTED: lachesin-like [*Hyaella azteca*] | *ncbi_113817215* | 3.8353 |
| 37 | *ncbi_113815885* | Transcription factor Sox-11-A [*Zootermopsis nevadensis*] | *ncbi_113815885* | 3.8098 |
| 38 | *Ttf2* | PREDICTED: transcription termination factor 2-like [*Hyaella azteca*] | *ncbi_113824440* | 3.8075 |
| 39 | *ncbi_113803783* | - | *ncbi_113803783* | 3.6866 |
| 40 | *ncbi_113803501* | - | *ncbi_113803501* | 3.5682 |
| 41 | *Gs2* | glutamine synthetase [*Penaeus monodon*] | *ncbi_113803894* | 3.5508 |
| 42 | *ncbi_113816639* | - | *ncbi_113816639* | 3.5447 |
| 43 | *zfh1* | PREDICTED: zinc finger protein 1 isoform X2 [*Aethina tumida*] | *ncbi_113824903* | 3.5015 |
| 44 | *ncbi_113800266* | PREDICTED: tumor protein p53-inducible protein 11-like [*Limulus polyphemus*] | *ncbi_113800266* | 3.4761 |
| 45 | *PRDX5* | peroxiredoxin-5 [*Palaemon carinicauda*] | *ncbi_113807833* | 3.3775 |
| 46 | *ncbi_113810290* | - | *ncbi_113810290* | 3.3480 |
| 47 | *Ras64B* | Ras protein [*Marsupenaeus japonicus*] | *ncbi_113809589* | 3.3452 |
| 48 | *ncbi_113812463* | - | *ncbi_113812463* | 3.2807 |
| 49 | *ncbi_113806506* | - | *ncbi_113806506* | 3.2742 |
| 50 | *ncbi_113810214* | - | *ncbi_113810214* | 3.2426 |
| 51 | *ncbi_113803496* | - | *ncbi_113803496* | 3.2234 |
| 52 | *Clca4a* | Calcium-activated chloride channel regulator 2 [*Daphnia magna*] | *ncbi_113800211* | 3.2105 |
| **53** | *Casp1* | PREDICTED: caspase-1 [*Monomorium pharaonis*] | ncbi_113800234 | 3.2027 |
| **54** | *ncbi_113805453* | PREDICTED: uncharacterized protein<br>LOC106655919 isoform X2 [*Trichogramma pretiosum*] | ncbi_113805453 | 3.2003 |
| **55** | *ncbi_113830025* | - | ncbi_113830025 | 3.1985 |
| **56** | *NPC2* | Niemann-Pick type C-2-like protein<br>[*Penaeus monodon*] | ncbi_113806913 | 3.1886 |
| **57** | *ncbi_113800422* | gill C-type lectin [*Marsupenaeus japonicus*] | ncbi_113800422 | 3.0434 |
| **58** | *Hr96* | Nuclear hormone receptor HR96 [*Daphnia magna*] | ncbi_113804121 | 2.9242 |
| **59** | *MP1* | serine protease 1 [*Litopenaeus vannamei*] | ncbi_113824294 | 2.9165 |
| **60** | *ncbi_113824399* | PREDICTED: uncharacterized protein<br>LOC100199872 [*Hydra vulgaris*] | ncbi_113824399 | 2.8101 |

**Table supplement 6.** Top 20 specifically expressed marker genes for prohemocyte 2 (PH2).

| # | Gene Name | Description | Gene ID | Log2FC |
| --- | --- | --- | --- | --- |
| 1 | *TGM1* | hemocyte transglutaminase [*Litopenaeus vannamei*] | ncbi_113823934 | 3.1294 |
| 2 | *ncbi_113805892* | OV-16 antigen [*Thelohanelus kitauei*] | ncbi_113805892 | 2.8076 |
| 3 | *ncbi_113823945* | hemocyte transglutaminase [*Litopenaeus vannamei*] | ncbi_113823945 | 2.7538 |
| 4 | *ncbi_113814237* | PREDICTED: neurogenic locus notch homolog protein 1-like [*Hyaella azteca*] | ncbi_113814237 | 2.7038 |
| 5 | *ncbi_113803465* | - | ncbi_113803465 | 2.4693 |
| 6 | *Igfbp7* | single insulin-like growth factor-binding domain protein-2 [*Cupiennius salei*] | ncbi_113811702 | 2.3611 |
| 7 | *ncbi_113808957* | PREDICTED: neurogenic locus notch homolog protein 1-like [*Hyaella azteca*] | ncbi_113808957 | 2.3587 |
| 8 | *PPAF2* | masquerade-like serine proteinase-like protein 3 [*Penaeus monodon*] | ncbi_113812814 | 2.2465 |
| 9 | *pyx* | PREDICTED: transient receptor potential cation channel subfamily A member 1-like [*Hyaella azteca*] | ncbi_113812900 | 2.2188 |
| 10 | *Grik2* | PREDICTED: glutamate receptor ionotropic, kainate 2 isoform X1 [*Polistes dominula*] | ncbi_113823802 | 2.1869 |
| 11 | *Cpa1* | PREDICTED: carboxypeptidase B-like [*Hyaella azteca*] | ncbi_113825742 | 2.0337 |
| 12 | *Rab27a* | Rab27A [*Macrobrachium rosenbergii*] | ncbi_113829827 | 2.0332 |
| 13 | *Ctrc* | prophenoloxidase activating enzyme [*Litopenaeus vannamei*] | ncbi_113819328 | 1.8991 |
| 14 | *ncbi_113800422* | gill C-type lectin [*Marsupenaeus japonicus*] | ncbi_113800422 | 1.8520 |
| 15 | *His2A* | PREDICTED: histone H2A-like [*Hyaella azteca*] | ncbi_113829109 | 1.8444 |
| 16 | *ncbi_113809022* | - | ncbi_113809022 | 1.8050 |
| 17 | *ncbi_113823668* | hypothetical protein [*Triatoma infestans*] | ncbi_113823668 | 1.6539 |
| 18 | *ncbi_113826594* | - | ncbi_113826594 | 1.6365 |
| 19 | *ncbi_113802865* | - | ncbi_113802865 | 1.6349 |
| 20 | *Mgst1* | microsomal prostaglandin E synthase [*Penaeus monodon*] | ncbi_113822637 | 1.6146 |

**Table supplement 7.** Top 20 specifically expressed marker genes for granulocyte 1 (GH1).

| # | Gene Name | Description | Gene ID | Log2FC |
| --- | --- | --- | --- | --- |
| 1 | *RGN* | PREDICTED: regucalcin-like [*Megachile rotundata*] | ncbi_113826969 | 2.5052 |
| 2 | *rgn* | PREDICTED: regucalcin-like isoform X2 [*Acromyrmex echinator*] | ncbi_113807911 | 2.1506 |
| 3 | *ncbi_113800677* | - | ncbi_113800677 | 2.1118 |
| 4 | *ncbi_113815774* | double WAP domain-containing protein [*Fenneropenaeus chinensis*] | ncbi_113815774 | 1.9894 |
| 5 | *ncbi_113802174* | stylicine 2, partial [*Litopenaeus stylirostris*] | ncbi_113802174 | 1.9801 |
| 6 | *PPAF2* | serine protease-like protein 3 [*Penaeus monodon*] | ncbi_113802490 | 1.9443 |
| 7 | *Arg2* | PREDICTED: arginase-1 isoform X2 [*Nanorana parkeri*] | ncbi_113811237 | 1.9291 |
| 8 | *ncbi_113813598* | single IB domain protein [*Litopenaeus vannamei*] | ncbi_113813598 | 1.7636 |
| 9 | *ncbi_113818447* | - | ncbi_113818447 | 1.7370 |
| 10 | *ncbi_113810419* | Juvenile hormone-inducible protein [*Culex quinquefasciatus*] | ncbi_113810419 | 1.7349 |
| 11 | *ncbi_113801793* | PREDICTED: kynurenine formamidase-like [*Hyalella azteca*] | ncbi_113801793 | 1.6970 |
| 12 | *ncbi_113821158* | crustin 2 [*Panulirus japonicus*] | ncbi_113821158 | 1.6816 |
| 13 | *ncbi_113802754* | crustacean hematopoietic factor-like protein [*Litopenaeus vannamei*] | ncbi_113802754 | 1.6322 |
| 14 | *Ces2c* | juvenile hormone esterase-like carboxylesterase 1 [*Eriocheir sinensis*] | ncbi_113812972 | 1.5684 |
| 15 | *PPAF2* | prophenoloxidase activating factor [*Litopenaeus vannamei*] | ncbi_113828564 | 1.5640 |
| 16 | *A2ML1* | alpha 2 macroglobulin [*Litopenaeus vannamei*] | ncbi_113826593 | 1.5613 |
| 17 | *SPE* | serine proteinase homologue, partial [*Marsupenaeus japonicus*] | ncbi_113828686 | 1.5507 |
| 18 | *PPO1* | prophenoloxidase 2 [*Litopenaeus vannamei*] | ncbi_113828743 | 1.5456 |
| 19 | *ncbi_113802179* | fortilin binding protein 1 [*Penaeus monodon*] | ncbi_113802179 | 1.5264 |
| 20 | *PPO3* | prophenoloxidase-1 [*Litopenaeus vannamei*] | ncbi_113827090 | 1.4719 |

**Table supplement 8.** Top 20 specifically expressed marker genes for monocytic hemocyte 1 (MH1).

| # | Gene Name | Description | Gene ID | Log2FC |
| --- | --- | --- | --- | --- |
| 1 | *ncbi_113809352* | single VWC domain protein 5 [*Litopenaeus vannamei*] | *ncbi_113809352* | 3.2121 |
| 2 | *ncbi_113823524* | - | *ncbi_113823524* | 3.0494 |
| 3 | *thbs4* | PREDICTED: cartilage oligomeric matrix protein-like isoform X2 [*Hyalella azteca*] | *ncbi_113823118* | 2.9563 |
| 4 | *ncbi_113800108* | - | *ncbi_113800108* | 2.8356 |
| 5 | *ncbi_113817695* | vascular endothelial growth factor 3 [*Litopenaeus vannamei*] | *ncbi_113817695* | 2.7241 |
| 6 | *ncbi_113800109* | - | *ncbi_113800109* | 2.4647 |
| 7 | *ncbi_113816490* | AAEL010263-PA [*Aedes aegypti*] | *ncbi_113816490* | 2.4272 |
| 8 | *ncbi_113823512* | - | *ncbi_113823512* | 2.3674 |
| 9 | *Nlrp3* | PREDICTED: NLR family CARD domain-containing protein 4-like [*Lingula anatina*] | *ncbi_113828560* | 2.2778 |
| 10 | *ncbi_113823500* | - | *ncbi_113823500* | 2.1004 |
| 11 | *ncbi_113816639* | - | *ncbi_113816639* | 2.0766 |
| 12 | *ncbi_113800107* | - | *ncbi_113800107* | 1.9934 |
| 13 | *ncbi_113830149* | hypothetical ferritin light-chain subunit, partial [*Rimicaris exoculata*] | *ncbi_113830149* | 1.8779 |
| 14 | *ncbi_113802107* | Ubiquitin [*Zootermopsis nevadensis*] | *ncbi_113802107* | 1.6898 |
| 15 | *nrv2* | sodium potassium-transporting ATPase subunit beta [*Litopenaeus vannamei*] | *ncbi_113810268* | 1.6152 |
| 16 | *Drice* | CAP-3 [*Litopenaeus vannamei*] | *ncbi_113805238* | 1.5990 |
| 17 | *Lyz1* | lysozyme [*Litopenaeus vannamei*] | *ncbi_113802295* | 1.5944 |
| 18 | *SULT1C4* | Sulfotransferase sult [*Daphnia magna*] | *ncbi_113827852* | 1.5719 |
| 19 | *ncbi_113820575* | PREDICTED: uncharacterized protein LOC108681463 [*Hyalella azteca*] | *ncbi_113820575* | 1.4612 |
| 20 | *ncbi_113823707* | - | *ncbi_113823707* | 1.4362 |

**Table supplement 9.** Source data file for qPCR.

| Region from FACS sorting | Biological replicates | Target genes | CT values |
| --- | --- | --- | --- |
| R1 | sample 1 | *EF-1α* | 29.94 |
| R1 | sample 2 | *EF-1α* | 31.10 |
| R1 | sample 3 | *EF-1α* | 29.83 |
| R1 | sample 4 | *EF-1α* | 31.68 |
| R1 | sample 5 | *EF-1α* | 27.97 |
| R2 | sample 1 | *EF-1α* | 25.79 |
| R2 | sample 2 | *EF-1α* | 27.28 |
| R2 | sample 3 | *EF-1α* | 27.65 |
| R2 | sample 4 | *EF-1α* | 30.84 |
| R2 | sample 5 | *EF-1α* | 26.47 |
| R1 | sample 1 | *CHIT1* | 34.37 |
| R1 | sample 2 | *CHIT1* | 32.58 |
| R1 | sample 3 | *CHIT* | 32.86 |
| R1 | sample 4 | *CHIT11* | 35.86 |
| R1 | sample 5 | *CHIT1* | 31.25 |
| R2 | sample 1 | *CHIT1* | 32.89 |
| R2 | sample 2 | *CHIT1* | 33.13 |
| R2 | sample 3 | *CHIT1* | 32.95 |
| R2 | sample 4 | *CHIT1* | 36.71 |
| R2 | sample 5 | *CHIT1* | 31.33 |
| R1 | sample 1 | *Lyz1* | 32.80 |
| R1 | sample 2 | *Lyz1* | 32.39 |
| R1 | sample 3 | *Lyz1* | 32.15 |
| R1 | sample 4 | *Lyz1* | 34.50 |
| R1 | sample 5 | *Lyz1* | 30.18 |
| R2 | sample 1 | *Lyz1* | 30.44 |
| R2 | sample 2 | *Lyz1* | 32.60 |
| R2 | sample 3 | *Lyz1* | 30.55 |
| R2 | sample 4 | *Lyz1* | 34.05 |
| R2 | sample 5 | *Lyz1* | 28.96 |
| R1 | sample 1 | *NAGA* | 33.79 |
| R1 | sample 2 | *NAGA* | 33.43 |
| R1 | sample 3 | *NAGA* | 32.58 |
| R1 | sample 4 | *NAGA* | 32.82 |
| R1 | sample 5 | *NAGA* | 30.63 |
| R2 | sample 1 | *NAGA* | 33.14 |
| R2 | sample 2 | *NAGA* | 33.55 |
| R2 | sample 3 | *NAGA* | 32.00 |
| R2 | sample 4 | *NAGA* | 33.42 |
| R2 | sample 5 | *NAGA* | 30.79 |

**Table supplement 10.**
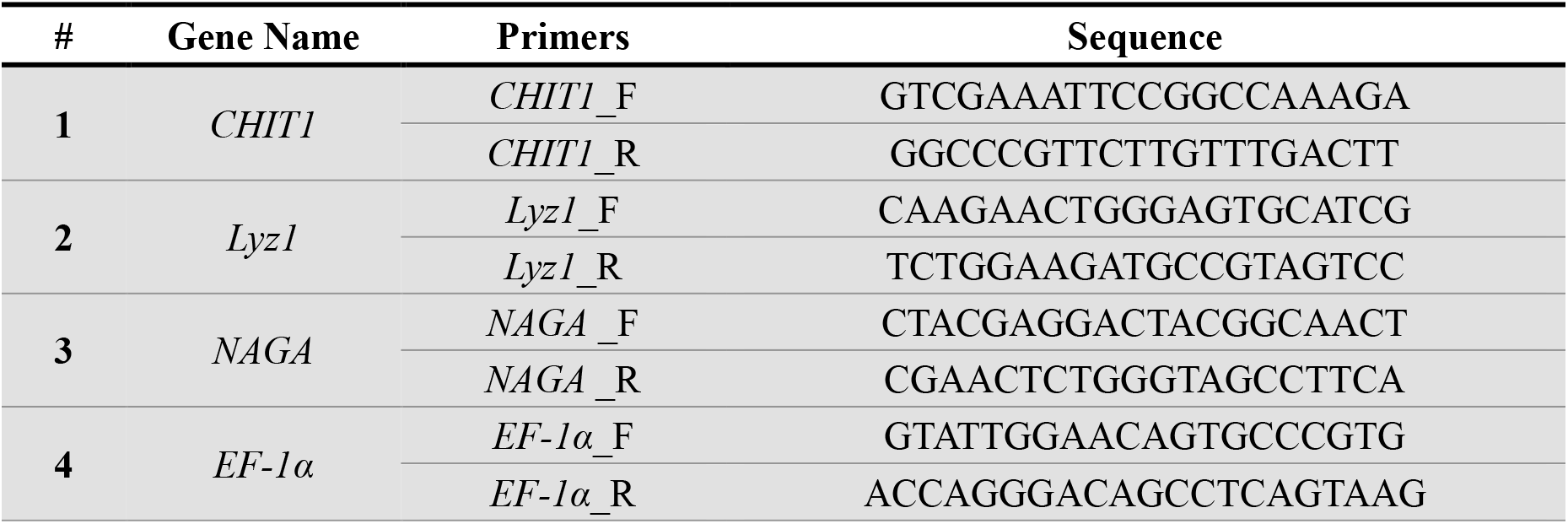
Primers used in this study.

## References

Balasubramanian, R., Bui, A., Xie, X., Deng, M., & Gan, L. (2014). Generation and characterization of Lhx9-GFPCreER(T2) knock-in mouse line. Genesis, 52(9), 827–832. doi:10.1002/dvg.22805

Bassler, K., Schulte-Schrepping, J., Warnat-Herresthal, S., Aschenbrenner, A. C., & Schultze, J. L. (2019). The Myeloid Cell Compartment-Cell by Cell. Annu Rev Immunol, 37, 269–293. doi:10.1146/annurev-immunol-042718-041728

Butler, A., Hoffman, P., Smibert, P., Papalexi, E., & Satija, R. (2018). Integrating single-cell transcriptomic data across different conditions, technologies, and species. Nat Biotechnol, 36(5), 411–420. doi:10.1038/nbt.4096

Cattenoz, P. B., Monticelli, S., Pavlidaki, A., & Giangrande, A. (2021). Toward a Consensus in the Repertoire of Hemocytes Identified in Drosophila. Front Cell Dev Biol, 9, 643712. doi:10.3389/fcell.2021.643712

Cho, B., Yoon, S. H., Lee, D., Koranteng, F., Tattikota, S. G., Cha, N., … Shim, J. (2020). Single-cell transcriptome maps of myeloid blood cell lineages in Drosophila. Nat Commun, 11(1), 4483. doi:10.1038/s41467-020-18135-y

Chung, N. C., & Storey, J. D. (2015). Statistical significance of variables driving systematic variation in high-dimensional data. Bioinformatics, 31(4), 545–554. doi:10.1093/bioinformatics/btu674

Friedman, A. D. (2002). Transcriptional regulation of granulocyte and monocyte development. Oncogene, 21(21), 3377–3390. doi:10.1038/sj.onc.1205324

Gong, Y., Wei, X., Sun, W., Ren, X., Chen, J., Aweya, J. J., … Li, S. (2021). Exosomal miR-224 contributes to hemolymph microbiota homeostasis during bacterial infection in crustacean. PLoS Pathog, 17(8), e1009837. doi:10.1371/journal.ppat.1009837

He, Y., Hara, H., & Nunez, G. (2016). Mechanism and Regulation of NLRP3 Inflammasome Activation. Trends Biochem Sci, 41(12), 1012–1021. doi:10.1016/j.tibs.2016.09.002

Huang, Z., Yang, P., & Wang, F. (2021). Shrimp Plasma CREG Is a Hemocyte Activation Factor. Front Immunol, 12, 707770. doi:10.3389/fimmu.2021.707770

Jo, Y. J., Lee, I. W., Jung, S. M., Kwon, J., Kim, N. H., & Namgoong, S. (2019). Spire localization via zinc finger-containing domain is crucial for the asymmetric division of mouse oocyte. FASEB J, 33(3), 4432–4447. doi:10.1096/fj.201801905R

Karlsson, M., Zhang, C., Mear, L., Zhong, W., Digre, A., Katona, B., Lindskog, C. (2021). A single-cell type transcriptomics map of human tissues. Sci Adv, 7(31). doi:10.1126/sciadv.abh2169

Kelly, L. M., Englmeier, U., Lafon, I., Sieweke, M. H., & Graf, T. (2000). MafB is an inducer of monocytic differentiation. EMBO J, 19(9), 1987–1997. doi:10.1093/emboj/19.9.1987

Koiwai, K., Koyama, T., Tsuda, S., Toyoda, A., Kikuchi, K., Suzuki, H., & Kawano, R. (2021). Single-cell RNA-seq analysis reveals penaeid shrimp hemocyte subpopulations and cell differentiation process. Elife, 10. doi:10.7554/eLife.66954

Kwon, H., Mohammed, M., Franzen, O., Ankarklev, J., & Smith, R. C. (2021). Single-cell analysis of mosquito hemocytes identifies signatures of immune cell subtypes and cell differentiation. Elife, 10. doi:10.7554/eLife.66192

Li, C., Yang, M. C., Hong, P. P., Zhao, X. F., & Wang, J. X. (2021). Metabolomic Profiles in the Intestine of Shrimp Infected by White Spot Syndrome Virus and Antiviral Function of the Metabolite Linoleic Acid in Shrimp. J Immunol, 206(9), 2075–2087. doi:10.4049/jimmunol.2001318

Li, H., Janssens, J., De Waegeneer, M., Kolluru, S. S., Davie, K., Gardeux, V., … Zinzen, R. P. (2022). Fly Cell Atlas: A single-nucleus transcriptomic atlas of the adult fruit fly. Science, 375(6584), eabk2432. doi:10.1126/science.abk2432

Li, T., Li, X., Zamani, A., Wang, W., Lee, C. N., Li, M., … Chen, Y. H. (2020). c-Rel Is a Myeloid Checkpoint for Cancer Immunotherapy. Nat Cancer, 1(5), 507–517. doi:10.1038/s43018-020-0061-3

Lin, X., & Soderhall, I. (2011). Crustacean hematopoiesis and the astakine cytokines. Blood, 117(24), 6417–6424. doi:10.1182/blood-2010-11-320614

Lun, A. T. L., Riesenfeld, S., Andrews, T., Dao, T. P., Gomes, T., participants in the 1st Human Cell Atlas, J., & Marioni, J. C. (2019). EmptyDrops: distinguishing cells from empty droplets in droplet-based single-cell RNA sequencing data. Genome Biol, 20(1), 63. doi:10.1186/s13059-019-1662-y

Luo, K., Chen, Y., & Wang, F. (2022). Shrimp Plasma MANF Works as an Invertebrate Anti- Inflammatory Factor via a Conserved Receptor Tyrosine Phosphatase. J Immunol. doi:10.4049/jimmunol.2100595

McNamara, J. C., & Faria, S. C. (2012). Evolution of osmoregulatory patterns and gill ion transport mechanisms in the decapod Crustacea: a review. J Comp Physiol B, 182(8), 997–1014. doi:10.1007/s00360-012-0665-8

Pan, C., & Fan, Y. (2016). Role of H1 linker histones in mammalian development and stem cell differentiation. Biochim Biophys Acta, 1859(3), 496–509. doi:10.1016/j.bbagrm.2015.12.002

Raddi, G., Barletta, A. B. F., Efremova, M., Ramirez, J. L., Cantera, R., Teichmann, S. A., Billker, O. (2020). Mosquito cellular immunity at single-cell resolution. Science, 369(6507), 1128–1132. doi:10.1126/science.abc0322

Short, M. L., Nickel, J., Schmitz, A., & Renkawitz, R. (1996). Lysozyme gene expression and regulation. EXS, 75, 243–257. doi:10.1007/978-3-0348-9225-4_13

Soderhall, I. (2016). Crustacean hematopoiesis. Dev Comp Immunol, 58, 129–141. doi:10.1016/j.dci.2015.12.009

Starck, J., Weiss-Gayet, M., Gonnet, C., Guyot, B., Vicat, J. M., & Morle, F. (2010). Inducible Fli-1 gene deletion in adult mice modifies several myeloid lineage commitment decisions and accelerates proliferation arrest and terminal erythrocytic differentiation. Blood, 116(23), 4795–4805. doi:10.1182/blood-2010-02-270405

Stuart, T., Butler, A., Hoffman, P., Hafemeister, C., Papalexi, E., Mauck, W. M., 3rd, Satija, R. (2019). Comprehensive Integration of Single-Cell Data. Cell, 177(7), 1888–1902 e1821. doi:10.1016/j.cell.2019.05.031

Sun, J., Wang, L., Yang, W., Li, Y., Jin, Y., Wang, L., & Song, L. (2021). A novel C-type lectin activates the complement cascade in the primitive oyster Crassostrea gigas. J Biol Chem, 297(6), 101352. doi:10.1016/j.jbc.2021.101352

Sun, M., Li, S., Zhang, X., Xiang, J., & Li, F. (2020). Isolation and transcriptome analysis of three subpopulations of shrimp hemocytes reveals the underlying mechanism of their immune functions. Dev Comp Immunol, 108, 103689. doi:10.1016/j.dci.2020.103689

Sun, R., Qiu, L., Yue, F., Wang, L., Liu, R., Zhou, Z., Song, L. (2013). Hemocytic immune responses triggered by CpG ODNs in shrimp Litopenaeus vannamei. Fish Shellfish Immunol, 34(1), 38–45. doi:10.1016/j.fsi.2012.09.016

Tao, M., Zhou, H., Luo, K., Lu, J., Zhang, Y., & Wang, F. (2019). Quantitative serum proteomics analyses reveal shrimp responses against WSSV infection. Dev Comp Immunol, 93, 89–92. doi:10.1016/j.dci.2019.01.003

Tassanakajon, A., Rimphanitchayakit, V., Visetnan, S., Amparyup, P., Somboonwiwat, K., Charoensapsri, W., & Tang, S. (2018). Shrimp humoral responses against pathogens: antimicrobial peptides and melanization. Dev Comp Immunol, 80, 81–93. doi:10.1016/j.dci.2017.05.009

Tattikota, S. G., Cho, B., Liu, Y., Hu, Y., Barrera, V., Steinbaugh, M. J., … Perrimon, N. (2020). A single-cell survey of Drosophila blood. Elife, 9. doi:10.7554/eLife.54818

Wang, G., Jin, S., Huang, W., Li, Y., Wang, J., Ling, X., … Li, X. (2021). LPS-induced macrophage HMGB1-loaded extracellular vesicles trigger hepatocyte pyroptosis by activating the NLRP3 inflammasome. Cell Death Discov, 7(1), 337. doi:10.1038/s41420-021-00729-0

Wang, G., Li, N., Zhang, L., Zhang, L., Zhang, Z., & Wang, Y. (2015). IGFBP7 promotes hemocyte proliferation in small abalone Haliotis diversicolor, proved by dsRNA and cap mRNA exposure. Gene, 571(1), 65–70. doi:10.1016/j.gene.2015.06.051

Xu, Z., Yang, Y., Sarath Babu, V., Chen, J., Li, F., Yang, M., … Qin, Z. (2021). The antibacterial activity of erythrocytes from Clarias fuscus associated with phagocytosis and respiratory burst generation. Fish Shellfish Immunol, 119, 96–104. doi:10.1016/j.fsi.2021.10.001

Yang, C. C., Lu, C. L., Chen, S., Liao, W. L., & Chen, S. N. (2015). Immune gene expression for diverse haemocytes derived from pacific white shrimp, Litopenaeus vannamei. Fish Shellfish Immunol, 44(1), 265–271. doi:10.1016/j.fsi.2015.02.001

Zhang, X., Yuan, J., Sun, Y., Li, S., Gao, Y., Yu, Y., … Xiang, J. (2019). Penaeid shrimp genome provides insights into benthic adaptation and frequent molting. Nat Commun, 10(1), 356. doi:10.1038/s41467-018-08197-4

Zheng, Z., Aweya, J. J., Bao, S., Yao, D., Li, S., Tran, N. T., … Zhang, Y. (2021). The Microbial Composition of Penaeid Shrimps’ Hepatopancreas Is Modulated by Hemocyanin. J Immunol, 207(11), 2733–2743. doi:10.4049/jimmunol.2100746

